# Estimating the rate of plasmid transfer with an adapted Luria–Delbrück fluctuation analysis

**DOI:** 10.1101/2021.01.06.425583

**Authors:** Olivia Kosterlitz, Adamaris Muñiz Tirado, Claire Wate, Clint Elg, Ivana Bozic, Eva M. Top, Benjamin Kerr

**Author notes:** **Author contributions**: O.K. and B.K. conceived of the presented ideas. B.K. and I.B. developed the theory. O.K. developed the simulations and experimental protocols. C.E. facilitated part of the simulations. O.K., A.M.T., and C.W. performed the experiments. O.K. and B.K. wrote the manuscript. E.M.T. and B.K. supervised the project. All authors discussed the results and contributed to the final manuscript. **Competing Interest Statement**: The authors declare no competing interests.

## Abstract

To increase our basic understanding of the ecology and evolution of conjugative plasmids, we need a reliable estimate of their rate of transfer between bacterial cells. However, accurate estimates of plasmid transfer have remained elusive due to biological and experimental complexity. Current methods to measure transfer rate can be confounded by many factors. A notable example involves plasmid transfer between different strains or species where the rate that one type of cell donates the plasmid is not equal to the rate at which the other cell type donates. Asymmetry in these rates has the potential to bias or constrain current transfer estimates, thereby limiting our capabilities for estimating transfer in microbial communities. Inspired by the classic fluctuation analysis of Luria and Delbrück, we develop a novel approach, the Luria-Delbrück method (‘LDM’), for estimating plasmid transfer rate. Our new approach embraces the stochasticity of conjugation departing from the current deterministic population dynamic methods. In addition, the LDM overcomes obstacles of traditional methods by not being affected by different growth and transfer rates for each population within the assay. Using stochastic simulations and experiments, we show that the LDM has high accuracy and precision for estimation of transfer rates compared to the most widely used methods, which can produce estimates that differ from the LDM estimate by orders of magnitude.

**Significance Statement:** Conjugative plasmids play significant roles in the ecology and evolution of microbial communities. Notably, antibiotic resistance genes are often encoded on conjugative plasmids. Thus, conjugation—the transfer of a plasmid copy from one cell to another—is a common way for antibiotic resistance to spread between important clinical pathogens. For both public health modeling and a basic understanding of microbial population biology, accurate estimates of this fundamental rate are of great consequence. We show that widely used methods can lead to biased estimates, deviating from true values by several orders of magnitude. Therefore, we developed a new approach, inspired by the classic fluctuation analysis of Luria and Delbrück, for accurately assessing the rate of plasmid conjugation under a variety of conditions.

## Introduction

A fundamental rule of heredity involves the passage of genes from parents to their offspring. Bacteria violate this rule of strict vertical inheritance by shuttling DNA between cells through horizontal gene transfer (1, 2). Often the genetic elements being shuttled are plasmids, extrachromosomal DNA molecules that can encode the machinery for their transfer (3). This plasmid transfer process is termed conjugation, in which a plasmid copy is moved from one cell to another upon direct contact. Additionally, plasmids replicate independently inside their host cell to produce multiple copies, which segregate into both offspring upon cell division. Therefore, conjugative plasmids are governed by two modes of inheritance: vertical and horizontal.

This horizontal mode of inheritance makes it possible for non-related cells to exchange genetic material, which includes members of different species (4). In fact, conjugation can occur across vast phylogenetic distances, such that the expansive gene repertoire in the “accessory” genome encoded on conjugative plasmids is shared among many microbial species (5). This ubiquitous genetic exchange reinforces the central role of conjugation in shaping the ecology and evolution of microbial communities (1, 3, 6). Notably, conjugation is a common mechanism facilitating the spread of antimicrobial resistance genes among bacteria and the emergence of multi-drug resistance in clinical pathogens (7–9). To understand how genes, including those of clinical relevance, move within complex bacterial communities, an accurate and precise measure of the rate of conjugation is of the utmost importance.

The basic approach to measure conjugation involves mixing plasmid-containing bacteria, called “donors”, with plasmid-free bacteria, called “recipients”. As the co-culture incubates, recipients acquire the plasmid from the donor through conjugation, and these transformed recipients are called “transconjugants”. Over the course of this “mating assay,” the densities of donors, recipients, and transconjugants are tracked over time (*D_t_, R_t_*, and *T_t_*, respectively) as the processes of population growth and plasmid transfer occur. To understand how such information is used to calculate the rate of conjugation, we consider an altered version of the foundational Levin *et al*. model (10). In this framework, populations grow exponentially, and recipients become transconjugants via conjugation when they interact with plasmid-bearing cells (i.e., donors or transconjugants). The densities of the populations are described by the following differential equations (the *t* subscript is dropped from the variables for notational convenience):

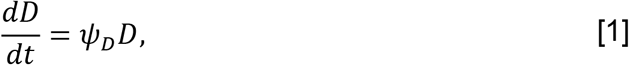

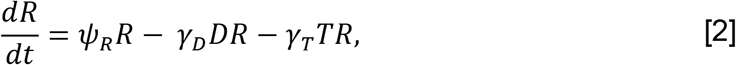

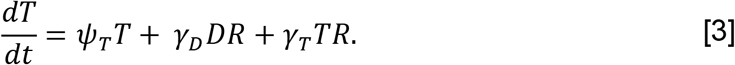

In equations [1]-[3], donors, recipients and transconjugants divide at a per-capita rate of *ψ_D_, ψ_R_*, and *ψ_T_*, respectively. The parameters *γ_D_* and *γ_T_* measure the rate at which a recipient cell acquires a plasmid per unit density of the donor and transconjugant, respectively. Thus, the *ψ* parameters are population growth rates and the *γ* parameters are conjugation rates (see Figure 1a). Assuming all the growth rates are equal (*ψ_D_* = *ψ_R_* = *ψ_T_* = *ψ*) and conjugation rates are equal (*γ_D_* = *γ_T_*=*γ*), Simonsen *et. al*. (11) provided an elegant solution to equations [1]-[3] to produce the following estimate for the conjugation rate from donors to recipients (hereafter termed the “donor conjugation rate”):

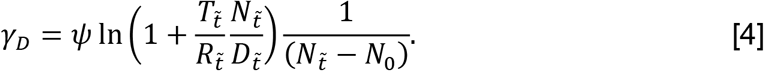

**Figure 1:**
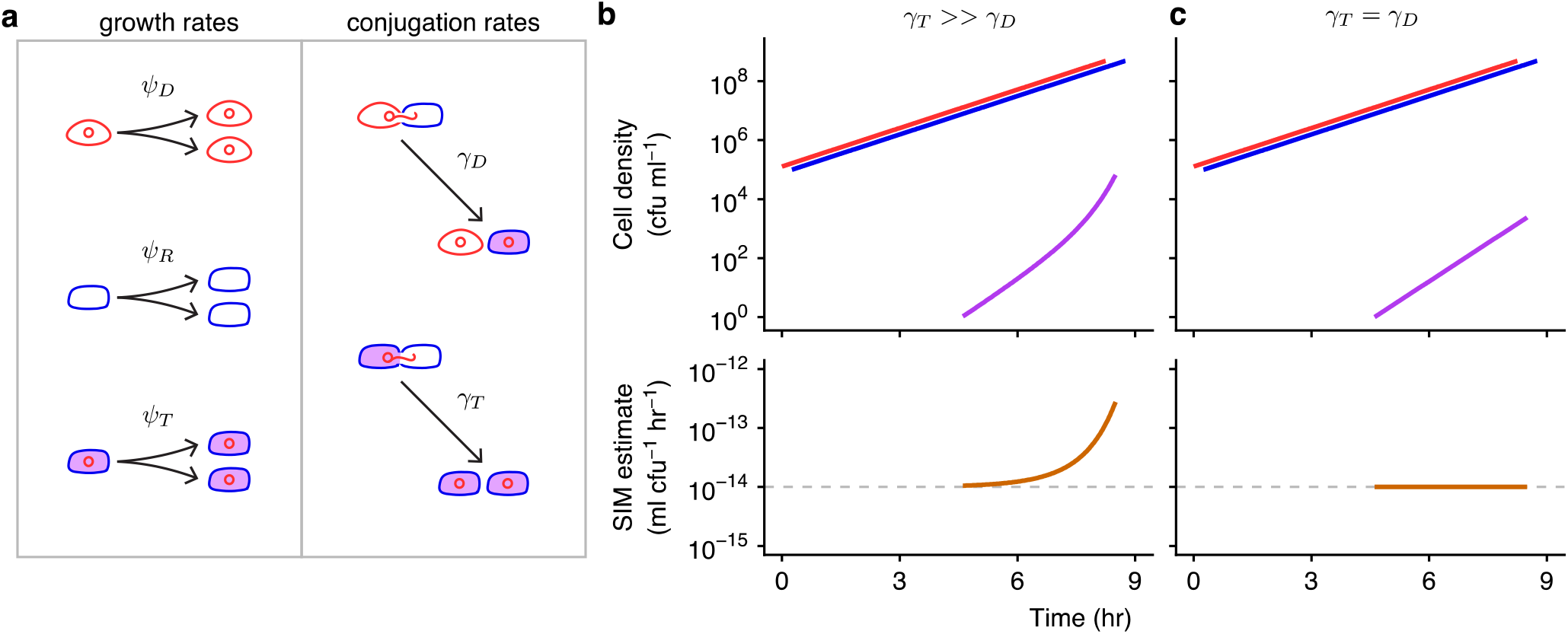
Basic model parameters and the effects of unequal conjugation rates on the SIM estimate. (a) In this schematic, the conjugative plasmid is a red circle, a donor is a red cell containing the plasmid, a recipient is a blue cell, and a transconjugant is indicated with a purple interior (a blue cell containing a red plasmid). The *ψ_D_, ψ_R_*, and *ψ_T_* parameters are donor, recipient, and transconjugant growth rates, respectively, illustrated by one cell dividing into two. The *γ_D_* and *γ_T_* parameters are donor and transconjugant conjugation rates, respectively, shown by conjugation events transforming recipients into transconjugants. (b) When the transconjugant conjugation rate (*γ_T_*) is higher than the donor conjugation rate (*γ_D_*), transconjugants exhibit super-exponential increase (purple curve) while donors and recipients increase exponentially (red and blue lines). The SIM estimate (orange line) increases over time, deviating from the actual donor conjugation rate (gray dashed line). (c) In contrast, when the conjugation rates are equal (*γ_T_* = *γ_D_*), the transconjugant increase is muted relative to part b (purple line). The SIM assumptions are met, and the estimate is constant and accurate over time (orange line). Equations [1]-[3] were used to produce the top graphs, with *D*_0_ = *R*_0_ = 10^5^, *T*_0_ = 0, *ψ_D_* = *ψ_R_* = *ψ_T_* = 1, *γ_D_* = 10^-14^, and either *γ_T_* = 10^-8^ (in part b) or *γ_T_* = 10^-14^ (in part c). The donor and recipient trajectories overlapped but were staggered for visibility. Equation [4] was used to produce the bottom graphs.

For a mating assay incubated for a fixed period (hereafter 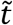), the initial and final density of all bacteria (*N*_0_ and 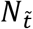, respectively), the final density of each cell population (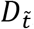, 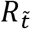, and 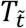), and the population growth rate (*ψ*) are sufficient for an estimate of the conjugation rate.

The Achilles heel of this estimate, as with others, is found in violations of its assumptions. For instance, we label equation [4] as the “Simonsen *et. al*. Identicality Method” estimate (SIM) for the donor conjugation rate because the underlying model assumes all strains are *identical* with regards to growth rates and conjugation rates. However, in natural microbial communities, this identicality assumption is misplaced, especially when the donors and recipients belong to different species. For instance, suppose that the rate of plasmid transfer within a species (i.e., from transconjugants to recipients, which we abbreviate as the “transconjugant conjugation rate”) is much higher than between species (i.e., from donors to recipients); that is, *γ_T_* ≫ *γ_D_* (Figure 1b). This elevated within-species conjugation rate (*γ_T_*) will increase the number of transconjugants and consequently inflate the SIM estimate for the cross-species conjugation rate (*γ_D_*) compared to a case where the conjugation rates are equal (*γ_T_* = *γ_D_*, Figure 1c). This Achilles heel is not specific to cross-species scenarios and can occur when estimating conjugation between any cells, including strains of the same species. One approach to minimize the resulting bias is to shorten the incubation time for the assay (12), as estimate bias tends to increase over time (e.g., Figure 1b). However, new problems can arise when using this approach, such as the transconjugant numbers becoming exceedingly low and thus difficult to accurately assess (13). Another approach was introduced by Huisman *et al*. (14), which squarely addressed the SIM identicality assumptions by developing a method to estimate donor conjugation rate when growth and transfer rates differ, thereby enlarging the set of systems amenable to estimation (see SI section 1 for full description of this and other approaches). Nonetheless, this new method can have difficulty with situations in which the donor conjugation rate (*γ_D_*) is substantially lower than the transconjugant rate (*γ_T_*), the example illustrated in Figure 1. Such differences have been reported in multi-species systems (15) and recently several studies have recognized the importance of evaluating the biology of plasmids in microbial communities (7, 16–18). Therefore, a method that provides an accurate estimate despite substantial inequalities in rate parameters is desirable.

Here we derive a novel estimate for conjugation rate, inspired by the Luria–Delbrück fluctuation experiment (19), by explicitly tracking transconjugant dynamics as a stochastic process (i.e., a continuous time branching process). Our method allows for unrestricted heterogeneity in growth rates and conjugation rates. Thus, our method fills a gap in the methodological toolkit by allowing unbiased estimation of conjugation rates in a wide variety of strains and species. We used stochastic simulations to validate our estimate and compare its accuracy and precision to other estimates. We developed a protocol for the laboratory by using microtiter plates to rapidly screen many donor-recipient co-cultures for the existence of transconjugants. In addition to its experimental tractability, our protocol circumvents problems that arise in the laboratory that can bias other approaches. Finally, we implemented our method in the laboratory and compared our estimate to the SIM estimate using a *Klebsiella pneumoniae* to *Escherichia coli* cross-species case study with an IncF conjugative plasmid.

## Results

### A new conjugation rate estimate inspired by the Luria–Delbrück approach

Previous methods to estimate the rate of conjugation have treated the rise of transconjugants as a deterministic process (i.e., non-random). However, conjugation is inherently a stochastic (i.e., random) process (20). Given that conjugation transforms the genetic state of a cell, we can form an analogy with mutation, which is also a stochastic process that transforms the genetic state of a cell. While mutation transforms a wild-type cell to a mutant, conjugation transforms a recipient cell to a transconjugant.

This analogy inspired us to revisit the way Luria and Delbrück handled the mutational process in their classic paper on the nature of bacterial mutation (19), outlined in Figure 2a-d. For this process, assume that the number of wild-type cells, *N_t_*, is expanding exponentially. Let the rate of mutant formation be given by *μ*. In Figure 2a, we see that the number of mutants in a growing population increases due to mutation events (highlighted purple cells) and due to faithful reproduction by mutants (non-highlighted purple cells). The rate at which mutants are generated (highlighted purple cells) is *μN_t_*, which grows as the number of wild-type cells increase (Figure 2b). However, the rate of transformation per wild-type cell is the mutation rate *μ*, which is constant (Figure 2c). Since mutations are random, parallel cultures will vary in the number of mutants depending on if and when mutation events occur. As seen in Figure 2d, for sufficiently small wild-type populations growing over sufficiently small periods, some replicate populations will not contain any mutant cell (gray shading) while other populations exhibit mutants (purple shading). Indeed, the cross-replicate fluctuation in the number of mutants was a critical component of the Luria-Delbrück experiment.

**Figure 2:**
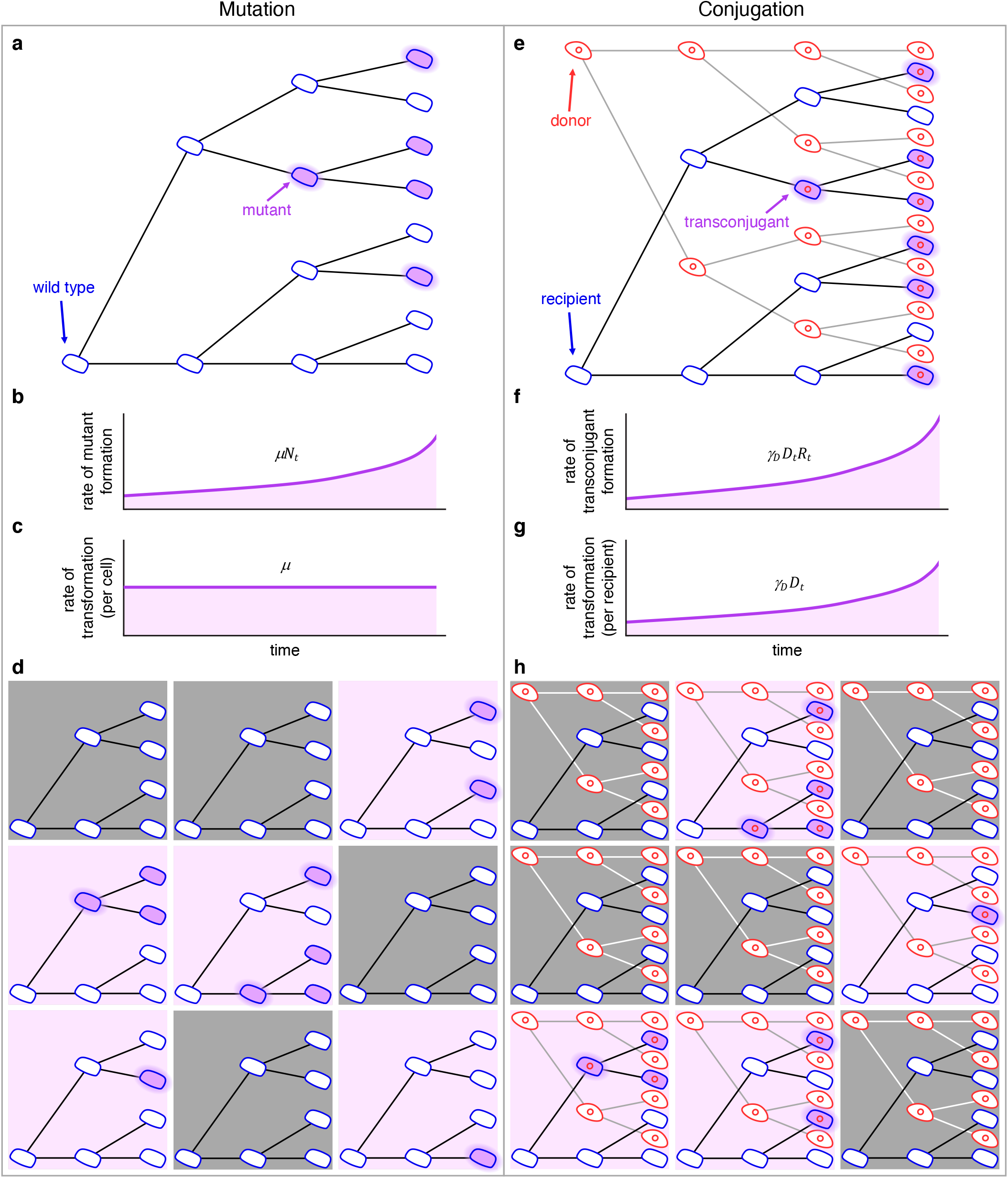
Schematic comparing the process of mutation (a-d) to the process of conjugation (e-h). (a) In a growing population of wild-type cells, mutants arise (highlighted purple cells) and reproduce (non-highlighted purple cells). (b) The rate at which mutants are generated grows as the number of wild-type cells increases (i.e., *μN_t_*). (c) The rate of transformation per wild-type cell is the mutation rate *μ*. (d) Wild-type cells growing in 9 separate populations where mutants arise in a portion of the populations (those with purple backgrounds) at different cell divisions. (e) In a growing population of donors and recipients, transconjugants arise (highlighted purple cells) and reproduce (non-highlighted purple cells). (f) The rate at which transconjugants are generated grows as the numbers of donors and recipients increase (i.e., *γ_D_D_t_R_t_*). (g) The rate of transformation per recipient cell grows as the number of donors increases (i.e., *γ_D_D_t_*) where *γ_D_* is the constant conjugation rate parameter. (h) Donor and recipient cells growing in 9 separate populations where transconjugants arise in a portion of the populations (purple backgrounds) at different points in time. For all panels, this is a conceptual figure, and the rates are inflated for illustration purposes.

To apply this strategy to estimate the conjugation rate, we can similarly think about an exponentially growing population of recipients (Figure 2e). But now there is another important cell population present (the donors). The transformation of a recipient is simply the generation of a transconjugant (highlighted purple cells) via conjugation with a donor. If we ignore conjugation from transconjugants for the moment, the rate at which transconjugants are generated is *γ_D_D_t_R_t_* (Figure 2f). In contrast to the mutation rate, the rate of transformation per cell is not a constant. Rather, this transformation rate per recipient is *γ_D_D_t_*, which grows with the donor population (Figure 2g). It is as if we are tracking a mutation process where the mutation rate is exponentially increasing. Yet the rate of transformation per recipient *and* donor is constant, which is the donor conjugation rate *γ_D_*. As with mutation, conjugation is random which results in a distribution in the number of transconjugants among parallel cultures depending on the time points at which transconjugants arise. As seen in Figure 2h, under certain conditions, some replicate populations will not contain any transconjugant cell (gray shading) while other populations will exhibit transconjugants (purple shading).

Using this analogy, here we describe a new approach for estimating conjugation rate which embraces conjugation as a stochastic process (20). Let the density of donors, recipients, and transconjugants in a well-mixed culture at time *t* be given by the variables *D_t_, R_t_*, and *T_t_*. In all that follows, we will assume that the culture is inoculated with donors and recipients, while transconjugants are initially absent (i.e., *D*_0_ > 0, *R*_0_ > 0, and *T*_0_ = 0). The donor and recipient populations grow according to the following standard exponential growth equations

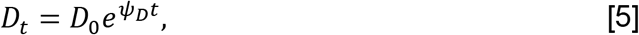

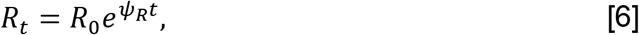

where *ψ_D_* and *ψ_R_* are the growth rates for donor and recipient cells, respectively. With equations [5] and [6], we are making a few assumptions, which also occur in some of the previous methods (SI Table 3). First, we assume the loss of recipient cells to transformation into transconjugants can be ignored. This assumption is acceptable because, for what follows, the rate of generation of transconjugants per recipient cell (as in Figure 2g, *γ_D_D_t_*) is very small relative to the per capita recipient growth rate (*ψ_R_*). Second, we assume that donors and recipients exhibit deterministic exponential growth. If the initial numbers of donors and recipients are not too small (i.e., *D*_0_ ≫ and *R*_0_ ≫ 0) and per capita growth remains constant over the period of interest, then this assumption is reasonable. We note that this assumption does not deny that cell division of donors and recipients is also a stochastic process, but given the large numbers of these cells, a deterministic approximation is appropriate.

On the other hand, the number of transconjugants over the period of interest can be quite small (starting from zero), motivating an explicit stochastic treatment (21). The population growth of transconjugants is modeled using a continuous-time stochastic process. The number of transconjugants, *T_t_*, is a random variable taking on non-negative integer values. In this section, we will assume the culture volume is 1 ml and thus the number of transconjugants is equivalent to the density of transconjugants (per ml). For a very small interval of time, Δ*t*, the current number of transconjugants will either increase by one or remain constant. The probabilities of each possibility are given as follows:

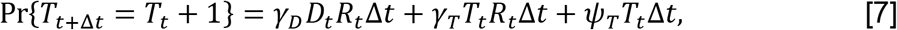

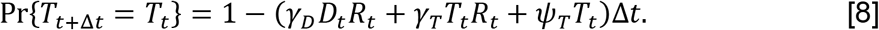

The three terms on the right-hand side of equation [7] illustrate the processes enabling the transconjugant population to increase. The first term gives the probability that a donor transforms a recipient into a transconjugant via conjugation. The second term gives the probability that a transconjugant transforms a recipient via conjugation. The third term measures the probability that a transconjugant cell divides. Equation [8] is simply the probability that none of these three processes occur.

Given the standard set-up of a mating assay, we focus on a situation where there are no transconjugants. Therefore, the only process that can change the number of transconjugants is conjugation of the plasmid from a donor to a recipient. Using equation [8] with *T_t_* = 0, we have

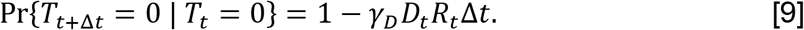

We let the probability that we have zero transconjugants at time *t* be denoted by *p*_0_(*t*) (i.e., *p*_0_(*t*) = Pr{ *T_t_* = 0}). In SI section 2, we derive the following expression for *p*_0_(*t*) at time 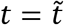:

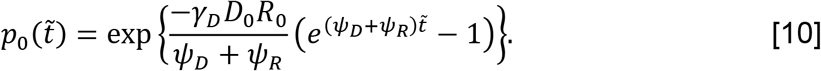

Solving equation [10] for *γ_D_* yields a new measure for the donor conjugation rate:

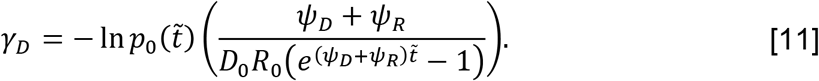

This expression is similar in form to the mutation rate derived by Luria and Delbrück in their classic paper on the nature of bacterial mutation (19), which is not a coincidence.

In SI section 3, we rederive the Luria-Delbrück result, which can be expressed as

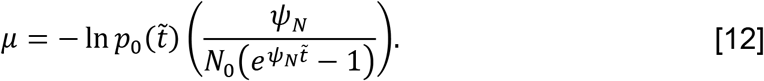

In the mutational process modeled by Luria and Delbrück, *N*_0_ is the initial wild-type population size, which grows exponentially at rate *ψ_N_*. For Luria and Delbrück, 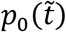 refers to the probability of zero mutants at time 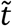 (as in a gray-shaded tree in Figure 2d), whereas 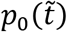 in the conjugation estimate refers to the probability of zero transconjugants (as in a gray-shaded tree in Figure 2h). Comparing equation [12] to equation [11], conjugation can be thought of as a mutation process with initial wild-type population size *D*_0_*R*_0_ that grows at rate *ψ_D_* + *ψ_R_*. We label the expression in equation [11] as the LDM estimate for donor conjugation rate, where LDM stands for “Luria-Delbrück Method” given the connection to their approach.

### The Luria-Delbrück method (LDM) has improved accuracy and precision

To explore the accuracy and precision of the LDM estimate and compare it to the SIM estimate (as well as other estimates, see SI section 4), we used the Gillespie algorithm to simulate the dynamics of a standard mating assay using equations [1]-[3] (Figure 3). Since the mating assay starts without transconjugants, a critical time point (hereafter *t*^*^) is marked by the creation of the first transconjugant cell due to the first conjugation event between a donor and a recipient. Before *t*^*^, the only events occurring are the cell divisions of donors and recipients (Figure 3a). After *t*^*^, all the event types described in Figure 1a can occur. Given that our simulation framework incorporates the stochastic nature of conjugation, *t*^*^ will vary among simulated mating assays. One stochastic run of the mating assay constitutes a simulation of the SIM approach. In the laboratory, the standard time point 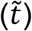 used for the SIM estimate is 24 hours, however, a truncated assay 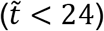 also produces a non-zero estimate of the conjugation rate as long as the incubation time is greater than *t*^*^ (the orange region of Figure 3b and c).

**Figure 3:**
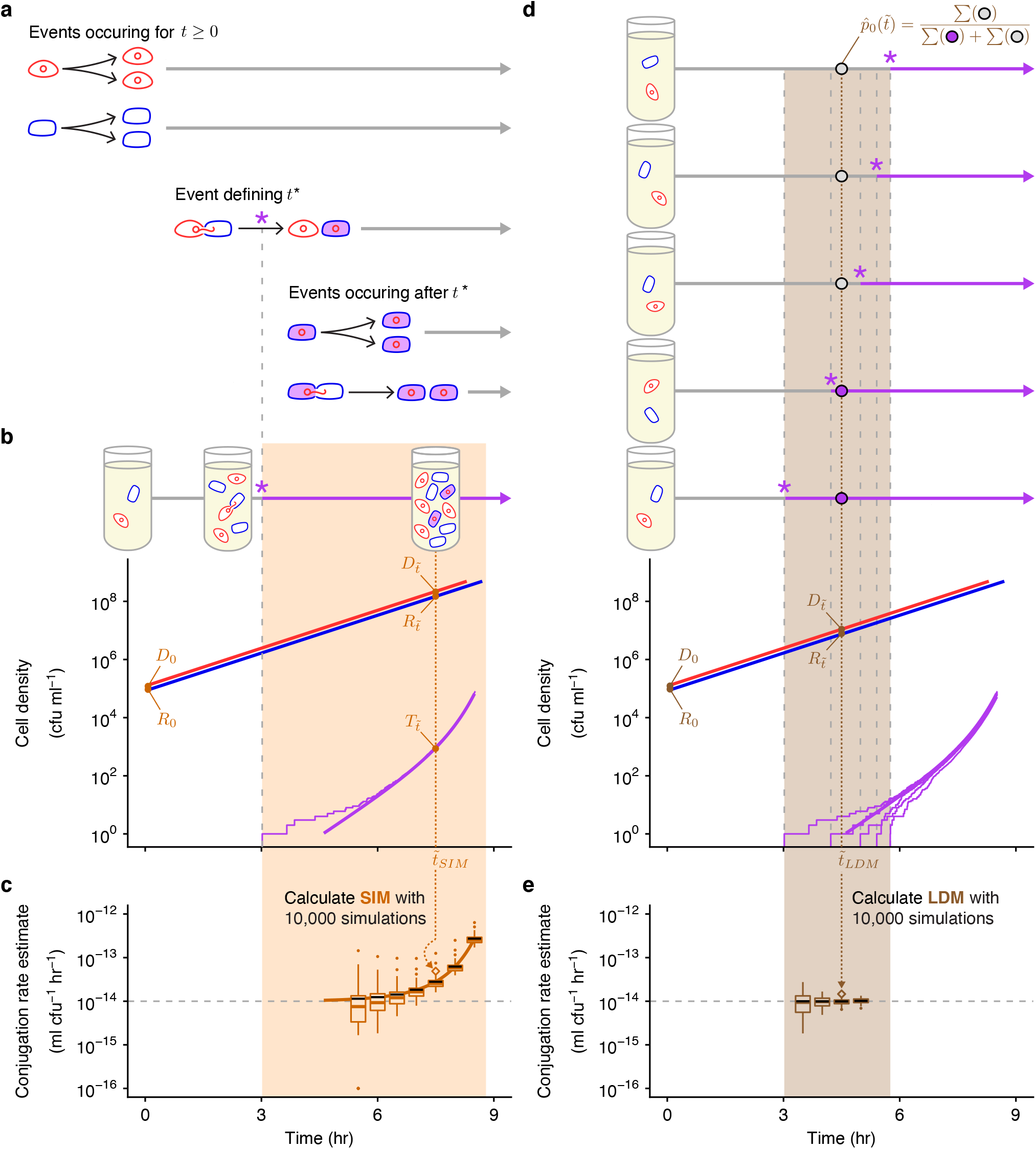
Overview of stochastic simulation framework and the effects of incubation time on estimating the conjugation rate. (a) The mating assay starts (*t* = 0) with donors and recipients and their populations increase over time. At a critical time (*t*^*^, marked by a purple asterisk), the first transconjugant cell is generated through a conjugation event between a donor and recipient. After *t*^*^, all possible growth and conjugation events can occur (including transconjugant division and conjugation). (b) A stochastic simulation of the equations [1]-[3] shows the donor, recipient, and transconjugant densities (red, blue, and purple thin trajectories, respectively) increasing over time. The deterministic numerical solution of the same equations and parameter settings from Figure 1b is shown for reference (thick lines). We note that for large densities, the stochastic and deterministic trajectories are closely aligned (i.e., the thick red and blue lines are overlaying their thin counterparts). After a specified incubation time (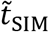, dotted orange line), we measure the densities of the three populations (orange 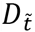, 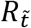, and 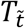), which can be used to calculate the (c) SIM estimate. (d) Multiple mating assays are needed for the LDM estimate. Here, five stochastic simulations are shown, which display variation in *t*^*^. At a specified incubation time (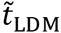, dotted brown line), we determine the number of assay cultures with transconjugants (purple circles, where for a relevant culture *i*, 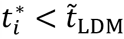) and without (gray circles, where for a relevant culture *j*, 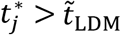). These numbers are used to calculate 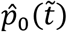, which, along with the donor and recipient densities (brown *D*_0_, *R*_0_, 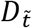 and 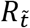) are used for the (e) LDM estimate. The SIM (part c) and LDM (part e) estimates are calculated for different incubation times, where the 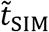 (part b) and 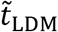 (part d) are indicated with orange and brown dotted arrows, respectively. The simulated trajectories in parts b and d would correspond to a single SIM or LDM estimate (the diamond points where the arrows terminate). The light orange and brown backgrounds indicate the range of incubation times giving a finite non-zero estimate of donor conjugation rate for the stochastic runs illustrated in parts b and d. In parts c and e, each box represents the estimate distribution using 10,000 simulations for a given 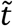, spanning from the 25th to 75th percentile. Given the log y-axis, the zero estimates are placed at the bottom of the y-axis range. The whiskers (i.e., vertical lines connected to the box) contain 1.5 times the interquartile range with the caveat that the whiskers were always constrained to the range of the data. The colored line in the box indicates the median. The solid black line indicates the mean. Parameter values are identical to Figure 1b and used throughout.

While the SIM estimate uses the density of transconjugants 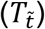, the LDM equation instead involves 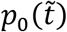, the probability that a population has no transconjugants at the end of the assay. A maximum likelihood estimate for this probability (hereafter 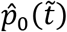) is obtained by calculating the fraction of populations (i.e., parallel simulations) that have no transconjugants at the specific incubation time 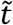 (top of Figure 3d). Thus, the range of time points to calculate the maximum likelihood estimate 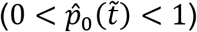 will be flanking the average *t*^*^ (the brown region of Figure 3d). Because the LDM estimate depends on the probabilistic *absence* of transconjugants, while the SIM estimate requires their *presence*, the range of incubation times for the LDM approach will be earlier than the SIM approach.

Even though there is a range of ‘valid’ incubation times, the accuracy of the SIM estimate can change over time as shown in Figure 3c (same case shown in Figure 1b). In this case, a key modeling assumption of the SIM approach was violated as the transconjugant conjugation rate was much higher than the donor conjugation rate (*γ_T_* ≫ *γ_D_*). Consequently, the SIM estimate of the donor conjugation rate was inflated compared to the true value by increasing amounts over time (Figure 3c). In contrast, the LDM estimate under the same scenario had high accuracy and precision over time (Figure 3e). We explored other parameter settings across various incubation times and the LDM estimate generally performed as well or better than other estimates (SI section 4).

To more systematically explore the effects of heterogeneous growth and conjugation rates on the accuracy and precision of estimating the donor conjugation rate (*γ_D_*), we ran sets of simulations sweeping through values of other parameters (*ψ_D_, ψ_R_, ψ_T_* and *γ_T_*). An illustrative example of heterogeneous growth occurs when plasmids confer costs or benefits on the fitness of their host. We simulated a range of growth-rate effects on plasmid-containing hosts from large plasmid costs (*ψ_D_* = *ψ_T_* ≪ *ψ_R_*) to large plasmid benefits (*ψ_D_* = *ψ_T_* ≫ *ψ_R_*). Relative to the SIM estimate, the LDM estimate had equivalent or higher accuracy and precision across all parameter settings (Figure 4a). To explore inequalities in conjugation rate more comprehensively, we simulated a range of transconjugant conjugation rates from relatively low (*γ_T_* ≪ *γ_D_*) to high (*γ_T_* ≫ *γ_D_*) values. Once again, the LDM estimate generally surpassed the SIM estimate across this range (Figure 4b). In SI section 4, we explore other parametric combinations along with model extensions, where, overall, the LDM outperformed the SIM approach and other estimates. Given the large number of simulations for these sweeps, we chose parameter values outside of experimentally obtained values reported in the literature, to reduce the computational burden of the Gillespie algorithm. However, the qualitative results were confirmed with a few simulations using parameter settings with more realistic values (SI section 4).

**Figure 4:**
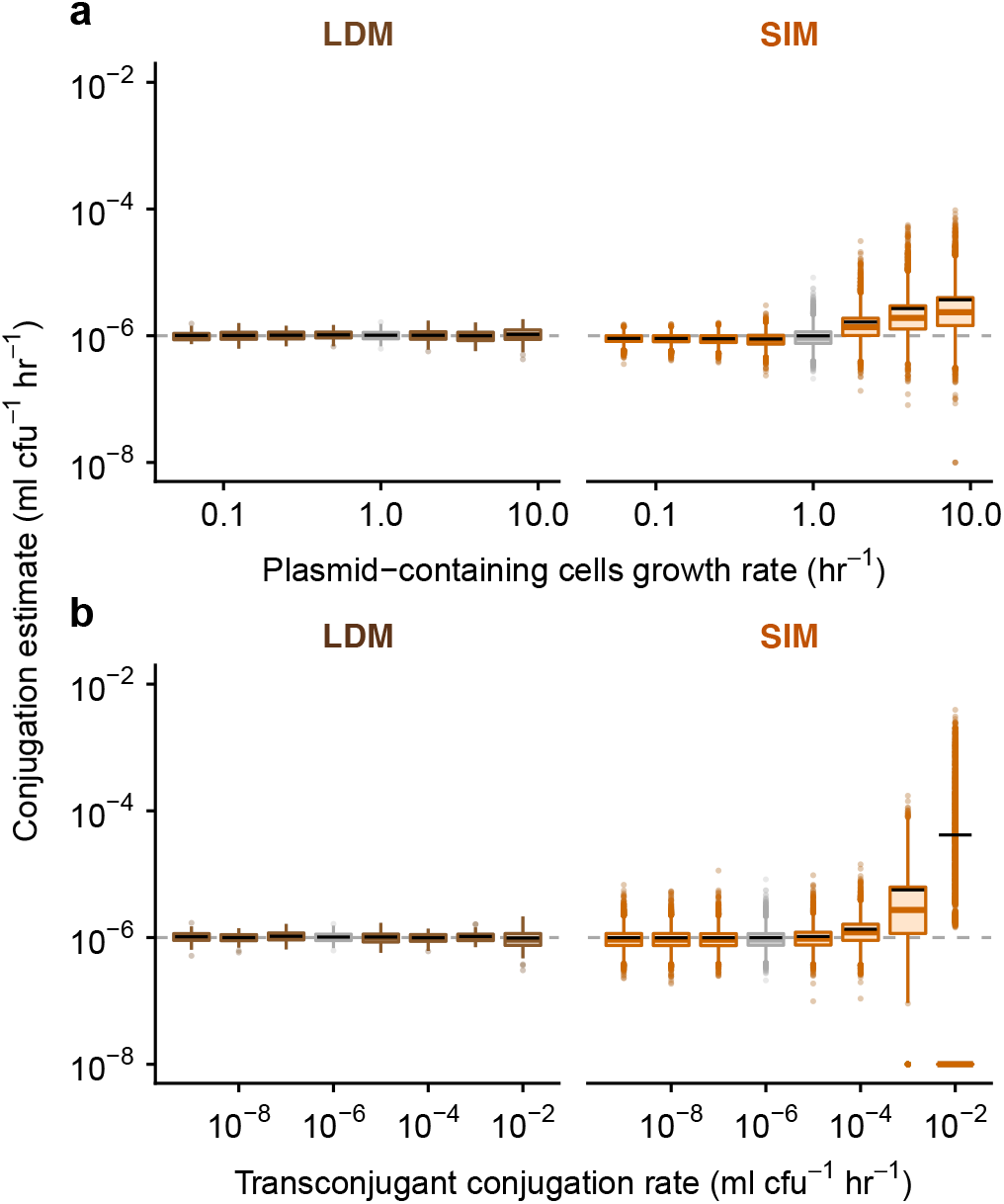
The effect of parametric heterogeneity on estimating conjugation rate. The Gillespie algorithm was used to simulate population dynamics. Donor conjugation rate for each parameter combination was estimated using 10,000 simulations (summarized using boxplots with the same graphical convention as in Figure 3). The gray dashed line indicates the true value for the donor conjugation rate (here, 10^-6^). The boxes in gray indicate the baseline parameter setting, and all colored boxes represent deviation of one or two parameters from baseline. The baseline parameter values were *ψ_D_* = *ψ_R_* = *ψ_T_* = 1 and *γ_D_* = *γ_T_* = 10^-6^. The dynamic variables were initialized with *D*_0_ = *R*_0_ = 10^2^ and *T*_0_ = 0. All incubation times are short but are specific to each parameter setting (see Materials and Methods). (a) Unequal growth rates were explored over a range of growth rates for the plasmid-bearing strains, namely *ψ_D_* = *ψ_T_* ∈ {0.0625, 0.125, 0.25, 0.5, 1, 2, 4, 8}. (b) Unequal conjugation rates were probed over a range of transconjugant conjugation rates, namely *γ_T_ ∈* {10^-9^, 10^-8^, 10^-7^, 10^-6^, 10^-5^, 10^-4^, 10^-3^, 10^-2^}. For the 10^-2^ transconjugant conjugation rate, many of the runs resulted in SIM estimates of zero; therefore, the median (colored line) and the box are placed at the bottom of the plot (given that the y-axis is on a log scale). The bulk of the data for this x-value is substantially lower than the mean SIM estimate (black line).

### New laboratory protocol to implement the LDM

We developed a general experimental procedure for estimating donor conjugation rate (*γ_D_*) using the LDM approach in the laboratory. The LDM protocol is tractable and can accommodate a wide variety of microbial species and conjugative plasmids by allowing for distinct growth and conjugation rates among donors, recipients, and transconjugants. The basic approach is to inoculate many donor-recipient co-cultures and then, at a time close to *t*^*^, add transconjugant-selecting medium (counterselection for donors and recipients) to determine the presence or absence of transconjugant cells in each co-culture.

In SI section 1, we rearrange equation [11] to provide an alternative form to highlight the quantities needed to conduct the LDM assay in the laboratory:

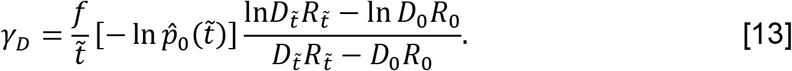

Similar to previous conjugation estimates, the LDM protocol requires measurement of initial and final densities of donors and recipients (*D*_0_, *R*_0_, 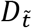, and 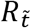). In addition, the LDM approach requires a fraction of parallel donor-recipient co-cultures to have no transconjugants at the specified incubation time 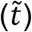, which is the maximum likelihood estimate 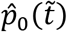. Lastly, there is a correction factor when the co-culture volume deviates from 1 ml; specifically, *f* is the reciprocal of the co-culture volume in ml (e.g., for a coculture volume of 100 μl, *f* = 1/0.1 = 10, SI section 5).

Before executing the LDM conjugation assay, an incubation time 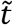 and initial density for the donors (*D*_0_) and the recipients (*R*_0_) needs to be chosen so that the probability that transconjugants form 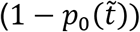 is not close to zero or one. We developed a short assay (SI section 6) for screening combinations of incubation time and initial densities to select a *target* incubation time 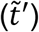 as well as *target* initial densities (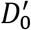 and 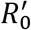) where 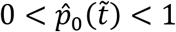. Note we add primes to indicate that these are ‘*targets*’ to distinguish *D*_0_, *R*_0_, and 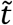 in equation [13] which will be gathered in the conjugation protocol itself. In addition, this pre-assay simultaneously verifies that the LDM modeling assumption of constant growth is satisfied. In our case, this pre-assay revealed several time-density combinations that could have been used. A useful pattern to note is that a higher donor conjugation rate will require shorter incubation times and lower initial densities compared to a lower rate.

For the LDM conjugation assay, we mix exponentially growing populations of donors and recipients, inoculate many co-cultures at the target initial densities in a 96 deep-well plate, and incubate in non-selective growth medium with the specific experimental culture volume (1/*f* of 1 ml) for the target incubation time (Figure 5). To estimate the initial densities (*D*_0_ and *R*_0_), three co-cultures at the start of the assay are diluted and plated on donor-selecting and recipient-selecting agar plates (Figure 5a). After the incubation time 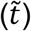, final densities (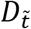 and 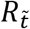) are also obtained by dilution-plating from the same co-cultures (Figure 5b). Liquid transconjugant-selecting medium is subsequently added to the remaining co-cultures (Figure 5c). After a long incubation in the transconjugant-selecting medium, there should be a mixture of turbid and non-turbid wells. A turbid well results from one or more transconjugant cells being present at time 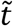 (when transconjugant-selecting medium was added). Therefore, a non-turbid well indicates the absence of transconjugant cells at 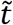, since the first conjugation event had not yet occurred (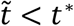, Figure 3), although see SI section 6. The proportion of non-turbid cultures is 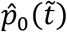 (Figure 5c). Unlike the traditional Luria–Delbrück method, no plating is required to obtain 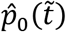. With the obtained densities (*D*_0_, *R*_0_, 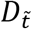, and 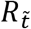), the incubation time 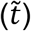, the proportion of transconjugant-free cultures 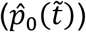, and the experimental culture volume correction (*f*), the LDM estimate for donor conjugation rate (*γ_D_*) can be calculated via equation [13].

**Figure 5:**
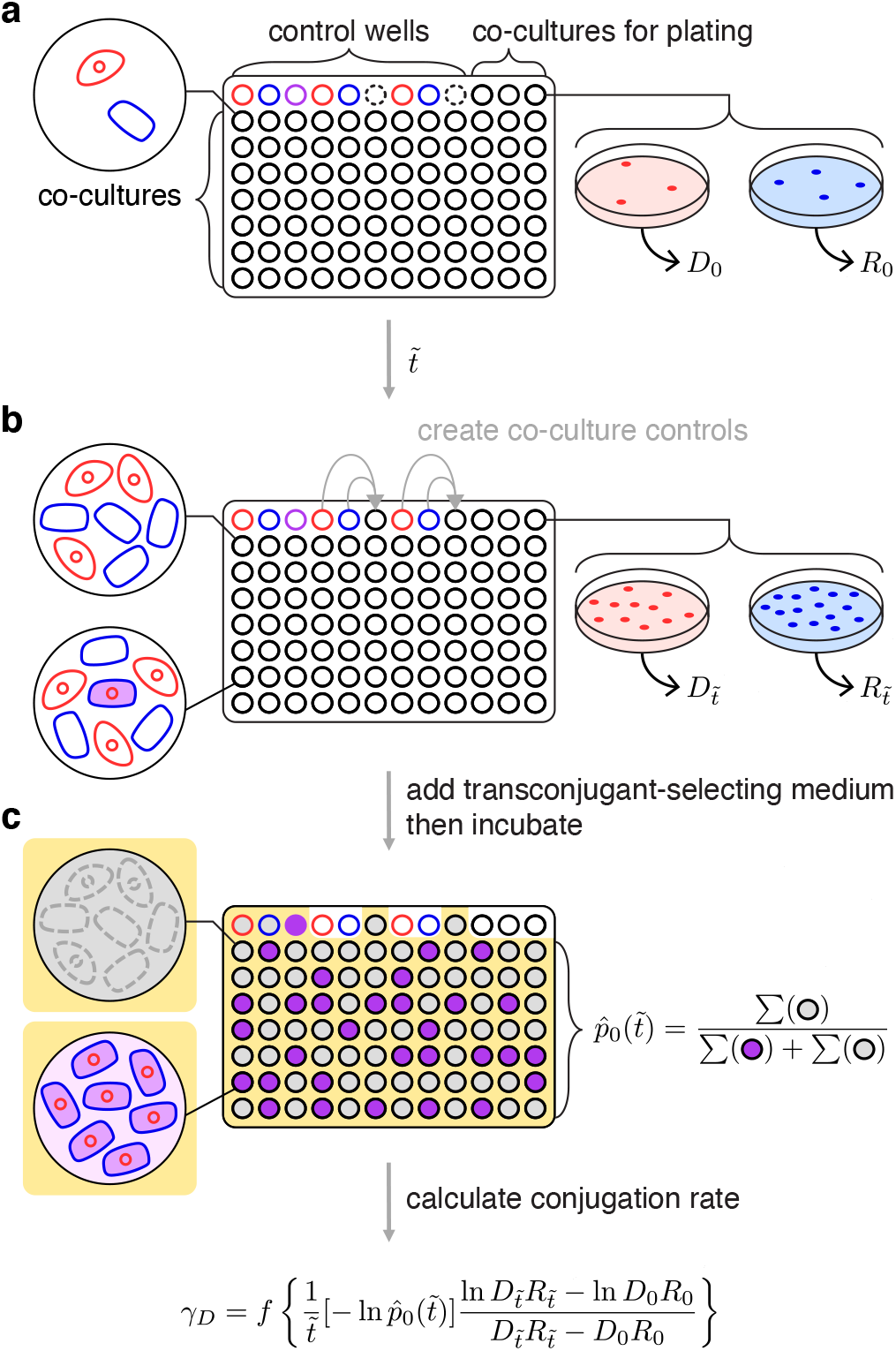
Overview for executing the LDM conjugation protocol. (a) The wells of a microtiter plate are inoculated with parallel co-cultures (black-bordered circles) at the target initial densities (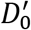 and 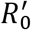). In addition, donor, recipient, and transconjugant monocultures serve as controls (red-, blue-, and purple-bordered wells, respectively). Three co-cultures (top-right) are sampled to determine the actual initial densities (*D*_0_ and *R*_0_). Note empty wells (dash-bordered circles) are used later in the assay. (b) After the incubation time 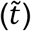, the same three co-cultures are sampled for final densities (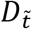 and 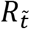). In addition, donor and recipient monocultures are mixed into the empty wells (indicated by grey arrows) to create co-culture controls to verify that diluting with transconjugant-selecting medium effectively prevents conjugation. (c) Subsequently, transconjugant-selecting medium is added to the microtiter plate (indicated by the yellow background) and incubated for a long period. The transconjugant-selecting medium should inhibit donor and recipient growth, leading to non-turbid (gray-filled) donor and recipient control wells, but a turbid (purple-filled) transconjugant control well. In addition, the transconjugant-selecting medium should prevent new conjugation events leading to non-turbid co-culture controls (gray-filled). Focusing on the wells inoculated with parallel co-cultures, the proportion of transconjugant-free (i.e., non-turbid, gray-filled) cultures is 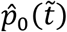. Using the actual incubation time 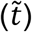, initial densities (*D*_0_ and *R*_0_), final densities (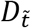 and 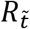), and the experimental culture volume correction (*f*), the LDM estimate of the donor conjugation rate (*γ_D_*) can be calculated. One microtiter plate yields one LDM estimate.

### Cross-species case study

To empirically test the performance of our assay and our modeling predictions, we initiated a cross-species mating assay between a donor, *Klebsiella pneumoniae* (hereafter ‘K’) with a conjugative IncF plasmid (hereafter ‘pF’), and a plasmid-free recipient, *Escherichia coli* (hereafter ‘E’). We denote the donor strain as K(pF), where the host species name is listed first and the plasmid inside the host is given in the parenthesis. E(Ø) denotes the plasmid-free recipient strain. We implemented the LDM and SIM protocols to estimate the cross-species conjugation rate in the laboratory.

The standard SIM protocol involves an incubation of 24 hours. For many bacterial species (including the ones explored here), an incubation time 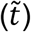 of 24 hours will lead to a violation of the assumption of constant growth rates from equations [1]-[3]. However, the original Simonsen et al. study did not actually assume constant growth rates (11). Their model permitted growth rate to vary as a function of resources, but additionally assumed that conjugation rate similarly varied. In other words, the ratio of growth and conjugation rates was assumed to remain constant (SI section 1). Under batch culture conditions, the population growth rates will drop as limiting resources are fully consumed (resulting in a stationary phase). As long as conjugation rates decrease with resources in a similar fashion and the parametric identicality assumptions hold, the SIM estimate can be used over a full-day incubation. We proceeded with the standard SIM protocol here.

Our LDM estimate of the cross-species conjugation rate was significantly lower than the standard SIM estimate, approximately three orders of magnitude (comparison A in Figure 6; t-test, p<0.05). This substantial incongruence could be due to a few possible factors. First, it is possible that the growth and conjugation rates do not change with nutrients in a functionally similar way. While we cannot rule out this possibility, it has been shown for IncF plasmids that both growth and conjugation drop as resources decline to low levels (10), consistent with SIM model assumptions. Second, our cell types have different growth rates (SI Figure 8), thus violating the SIM assumptions. While simulations show there is an effect of these inequalities, the effect size is insufficient to explain the observed difference in comparison A (SI section 4). Lastly, it is possible that the within-species conjugation, between the E(pF) transconjugants and E(Ø) recipients, occurs at a substantially higher rate than the cross-species conjugation, between the K(pF) donors and E(Ø) recipients. Our simulations show that this kind of difference in conjugation rates can lead to notable inflation of the SIM estimate, and there is evidence that within-species conjugation rates can be markedly elevated over cross-species rates (15, 22). Thus, this last possibility warranted further investigation.

**Figure 6:**
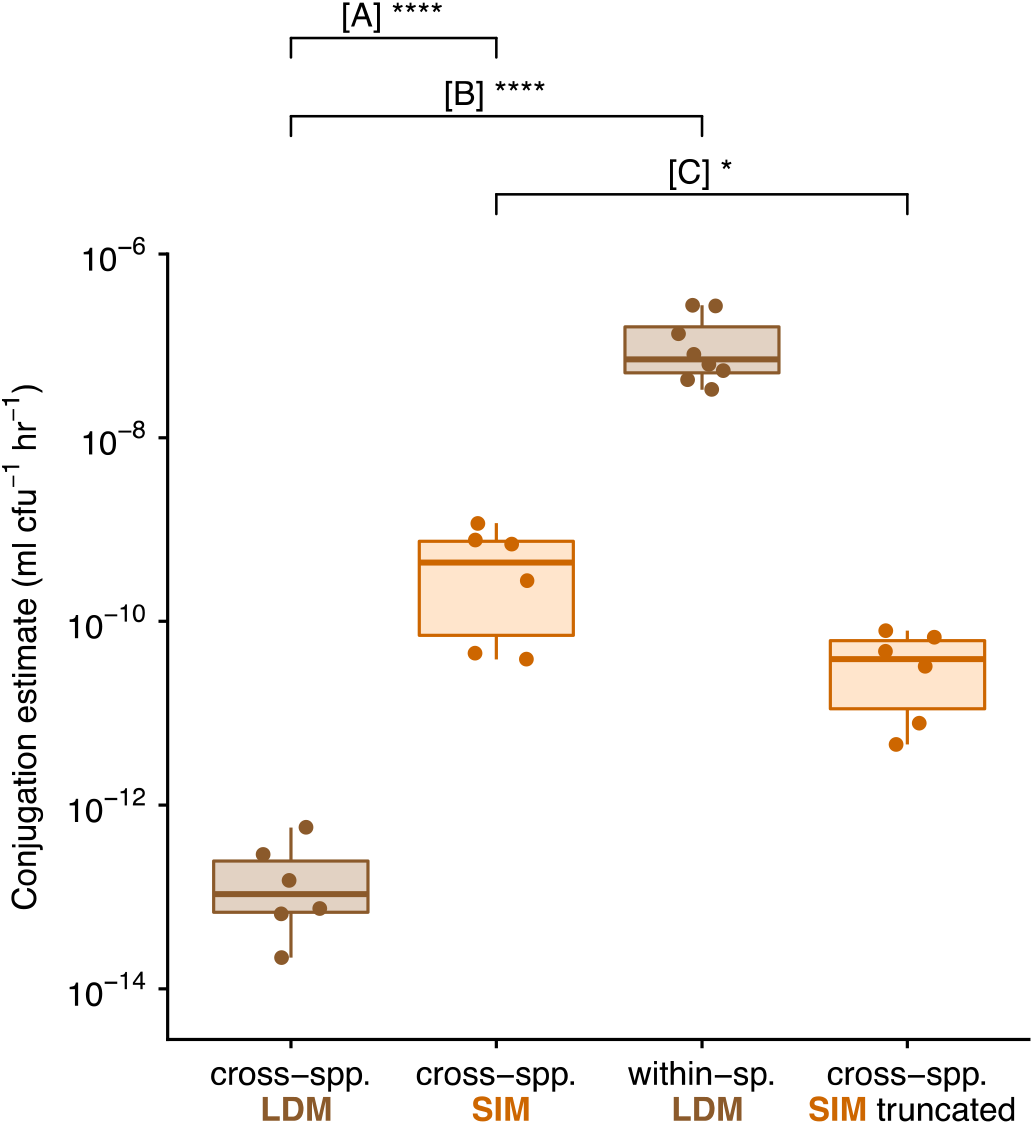
Experimental estimates for cross-species and within-species conjugation rates. Each box summarizes six replicate estimates by the LDM, SIM or truncated SIM approach, where each data point corresponds to a replicate. We note each of these estimates involved a correction (see Materials and Methods), but the same patterns hold for uncorrected values. [A] compares the LDM and standard SIM approach for a cross-species mating (between *K. pneumoniae* and *E. coli*). [B] compares the cross- and within-species mating using the LDM approach. [C] compares the standard and truncated SIM approach for a cross-species mating. The asterisks indicate statistical significance by a t-test (one, three and four asterisks convey p-values in the following ranges: 0.01 < p < 0.05, 0.0001 < p < 0.001 and p < 0.0001, respectively).

Next, we performed the within-species mating between *E. coli* strains. The LDM estimate for within-species conjugation rate (within *E. coli*) was higher than the cross-species LDM estimate by almost six orders of magnitude (comparison B in Figure 6; t-test, p<0.001), a difference that could explain the inflated SIM estimate. To further explore this explanation, we performed an additional cross-species SIM experiment with a shorter incubation time. In Figure 3c, as the incubation time was shortened, the SIM estimate approached the LDM estimate of the donor conjugation rate. Running the SIM protocol with a truncated incubation period (5 hours) resulted in a significantly lower cross-species conjugation rate estimate relative to the standard SIM estimate (comparison C in Figure 6; t-test, p<0.05), a result consistent with the pattern predicted under heterogeneous conjugation rates.

## Discussion

Conjugation is one of the primary modes of horizontal gene transfer in bacteria, facilitating the movement of genetic material between non-related neighboring cells. In microbial communities, conjugation can lead to the dissemination of genes among distantly related species. Since these genes are often of adaptive significance (e.g., antibiotic resistance), a comprehensive understanding of microbial evolution requires a full account of the process of conjugation. One of the most fundamental aspects of this process is the rate at which it occurs. Here we have presented a new method for estimating the rate of plasmid conjugative transfer from a donor cell to a recipient cell. We derived our LDM estimate using a mathematical approach that captures the stochastic process of conjugation, which was inspired by the method Luria and Delbrück applied to the process of mutation (19). We explored the connection between mutation and conjugation further in SI section 7. Our new method departs from the mathematical approach for other conjugation rate estimates, which assume underlying deterministic frameworks guiding the dynamics of transconjugants (10, 11, 14). Beyond the incorporation of stochasticity, the model and derivation behind the LDM estimate relaxes assumptions that constrain former approaches, which makes calculating conjugation rates accessible to a wide range of experimenters that use different plasmid-donor-recipient combinations.

### The LDM approach has improved accuracy

The most widely used approaches to estimate conjugation rate are derived from the Levin *et al*. model (SI section 1) which assumes that all strains grow and conjugate at the same rate (*ψ_D_* = *ψ_R_* = *ψ_T_* and *γ_D_* = *γ_T_*). These assumptions and constraints are problematic because bacterial growth and conjugation can and do vary (23, 24). Specifically, donors and recipients are often different taxa and contain chromosomal differences that translate to growth or conjugation rate differences (*ψ_D_* ≠ *ψ_R_* or *γ_D_* ≠ *γ_T_*). Additionally, plasmid carriage can change growth rate substantially (25) and therefore recipients can grow differently from donors (*ψ_R_* ≠ *ψ_D_*) or transconjugants (*ψ_R_* ≠ *ψ_T_*). In microbial communities, heterogeneous rates of growth and conjugation are the rule and not the exception. Therefore, a general estimation approach should be robust to this heterogeneity. While the estimates of popular approaches are insensitive to certain forms of heterogeneity, they can also be inaccurate under other forms. In contrast, the LDM estimate remains accurate across a broad range of heterogeneities.

A recent approach by Huisman *et al*. (14) relaxed the assumption of parametric homogeneity, yielding useful revisions to the SIM approach. However, when transconjugants exhibit much larger rates of plasmid transfer than the donors (*γ_T_* ≫ *γ_D_*), this new method can become inapplicable. Unfortunately, this kind of difference in conjugation rates is likely not uncommon in microbial communities (15, 26). Indeed, a mating assay involving two species can be thought of as a miniaturized microbial community where cross-species conjugation (between donors and recipients) and within-species conjugation (between transconjugants and recipients) both occur. Both previous work (15, 26) and experimental data from this study (Figure 6) demonstrate that the transconjugant (within-species) conjugation rate can be significantly higher than the donor (cross-species) rate. In addition, a similar difference in conjugation rates can arise from transitory de-repression, a molecular mechanism encoded on the conjugative plasmid that temporarily elevates the conjugation rate of a newly formed transconjugant (10, 27). The LDM approach is robust to these differences because it focuses on the creation of the first transconjugant (an event that must be between a donor and recipient) and ignores subsequent transconjugant dynamics (which is affected by transconjugant transfer). The LDM method produces an accurate estimate for donor conjugation rate in systems with unequal conjugation rates, whether the differences are taxonomic or molecular in origin.

### The LDM approach has improved precision

In addition to improved accuracy, the LDM estimate has advantages in terms of precision. Since conjugation is a stochastic process, the number of transconjugants at any given time is a random variable with a certain distribution. Therefore, estimates that rely on the number of transconjugants (which includes nearly all available methods) or the probability of their absence (the LDM approach) will also fall into distributions. Even in cases where the mean (first moment of the distribution) is close to the actual conjugation rate, the variance (second central moment) may differ among estimates. For the number of parallel co-cultures in our protocol, the LDM estimate had smaller variance compared to other estimates, even under parameter settings where different estimates shared similar accuracy (e.g., Figure 4). This greater precision likely originates from the difference in the distribution of the number of transconjugants 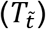 and the distribution of the probability of transconjugant absence 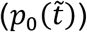, something we explore analytically in SI section 8. Beyond the mean and variance, other features of these distributions (i.e., higher moments) may also be important. For certain parameter settings, the estimates relying on transconjugant numbers were asymmetric (the third moment was non-zero). In such cases, a small number of replicate estimates could lead to bias (SI section 4). Typically, a small number of conjugation assays is standard; thus, the general position and shape of these estimate distributions may matter. Over the portion of parameter space that we explored, the LDM distribution facilitated accurate and precise estimates through its position (a mean reflecting the true value) and its shape (a small variance and a low skew).

### The LDM approach has implementation advantages

As discussed above, violations of modeling assumptions can lead to significant bias when estimating conjugation rate. Therefore, when implementing a conjugation protocol, the degree to which the experimental system satisfies the relevant assumptions is of prime importance. The most straightforward way to deal with this issue is to experimentally confirm that assumptions hold. For instance, the model underlying the LDM estimate assumes that growth rates of each cell type remain constant throughout the assay. This verification is part of the LDM protocol (see Materials and Methods and SI section 6). We emphasize that confirming the satisfaction of an assumption for one experimental system does not guarantee that the assumption holds for other systems. For example, the model underlying the SIM estimate assumes that growth and conjugation rates respond in a functionally similar way to changes in resources. While this assumption was verified for the IncF plasmid used in the original SIM study (10), other plasmid systems will readily violate it (e.g., some IncP plasmids conjugate during stationary phase after growth has stopped (28)), which can lead to bias in the estimate (SI section 4). Some approaches do not experimentally verify modeling assumptions as part of their corresponding protocol, but rather rely on simulated sensitivity analyses showing violations have little to no effect on the estimate (11, 14). For instance, the SIM estimate is robust to relatively small differences in growth rates or conjugation rates (11). Overall, for any conjugation rate estimate, either the underlying assumptions should be validated for the focal experimental system, or a rationale offered for why certain violations by the focal system will not significantly bias the estimate.

Given recent interest in the impacts of model assumption violations on conjugation rate estimates (14, 29), there has been a matching interest in altering conjugation protocols such that bias is minimized when violations apply. A common procedural adjustment involves shortening the incubation period of the mating assay because the bias resulting from modeling violations can increase over time (12, 13). For instance, when the transconjugant transfer rate is much higher than the donor rate, shortening the incubation time can mitigate some of the inaccuracies in the SIM estimate (Figure 3 and 6). However, there are a few caveats to this adjustment for estimates that rely on transconjugant density (which includes all common approaches, but not the LDM). First, as the incubation time decreases, the benefits in estimate accuracy come at the expense of costs in estimate precision. Specifically, variation in the timing of the first transconjugant cell appearance (*t*^*^ in Figure 3) has a greater impact on estimate variance with earlier incubation times. In part because the LDM approach does not rely on a measurement of transconjugant density, the LDM estimate remains both accurate and precise across various incubation times. Second, as incubation time decreases the transconjugant population can become extremely small and therefore technical problems with measuring an accurate transconjugant density through plating can arise (13). For instance, when the transconjugants are rare in the mating culture, the low dilution factor required for selective agar plating for transconjugants ensures a very high density of donors and recipients are simultaneously plated. Before complete inhibition of the donors and recipients by the transconjugant-selecting medium, conjugation events on the plate can generate additional transconjugants inflating the conjugation rate estimate (7, 13, 30, 31). Recently, a spectrophotometric technique was introduced to avoid selective plating altogether, which addresses this second caveat (13), but not the first. Notably, neither of these two caveats apply to the LDM approach because a binary output (turbid or non-turbid cultures) is used in lieu of measuring transconjugant density. Overall, the LDM protocol is both experimentally streamlined and insensitive to factors that can confound other approaches.

### The LDM approach is broadly applicable

In this paper, we have highlighted the possibility that the rate of conjugation may change (substantially) with the identity of the plasmid-bearing cell (32–34). For instance, as a plasmid moves from the original donor strain to the recipient background (forming a transconjugant), the transfer rate can change (i.e., *γ_D_* ≠ *γ_T_*). However, the conjugation rate changes with much more than just the identity of the cell holding the plasmid. The rate of transfer can additionally depend on the identity of the recipient as well as environmental conditions (e.g., level of nutrients, presence of antibiotics, etc.) (35). Thus, there is no *single* conjugation rate “belonging to” a plasmid-bearing strain. Some previous conjugation estimate methods (e.g., the SIM approach) build *conditionality* into their underlying model (e.g., conjugation rate changes dynamically with limiting resources). In such a case, the estimate is for a parameter (e.g., maximum conjugation rate) of the functional response, although additional assumptions about the functional response (e.g., conjugation and growth change proportionally) may introduce new methodological limitations. Our LDM approach is meant to be a conditional “snapshot,” where the conjugation rate depends on conditions of the protocol and the strains used. It is entirely possible to run the LDM approach under different conditions (e.g., changing nutrients) and assess the effect of environmental factors on transfer rate. The donor conjugation rate can be calculated under any condition as long as strain growth rates are constant over the protocol. But the distinguishing feature that gives the LDM method relative breadth of application is that it is robust to a form of conditionality that is tied to the mating assay itself. Specifically, because transconjugants are formed during a mating assay and, like donors, can deliver the plasmid to additional recipients, a form of rate conditionality is an unavoidable possibility for any protocol employing a mating assay. As we have shown (Figure 1, 3, 4, and 6), a difference in transfer rate between donors and transconjugants can make popular estimates inaccurate. However, by focusing on the first transconjugant formed (which only involves the donor and recipient, Figure 6), the LDM sidesteps this conditionality altogether, allowing an unbiased estimate of donor conjugation rate under a user-defined environment.

In conclusion, the LDM offers new possibilities for measuring the conjugation rate for many types of plasmids, species, and environmental conditions. We have presented evidence that supports our method being more accurate and precise than other widely used approaches. Importantly, the LDM eliminates bias caused by relatively high transconjugant conjugation rates, which is not unlikely when the donor and recipient belong to different species. We experimentally explored a case where the transconjugant transfer rate was dramatically higher than the donor rate and found that a standard estimate could inflate the conjugation rate (Figure 6). More generally, violations of model assumptions, intrinsic stochasticity, and implementation constraints can cause problems for currently available approaches. However, an adjustment of the approach Luria and Delbrück used to explore and estimate mutation over 75 years ago can address many of these issues. This new approach greatly expands the ability of experimentalists to accurately measure conjugation rates under the diverse conditions found in natural microbial communities.

## Materials and Methods

More detailed information for the mathematical models, simulations, and experiments are provided in the Supplementary Information.

### Bacterial Strains, Media, and Culture Conditions

Donor strains included two Enterobacteriaceae species: *Escherichia coli* K-12 BW25113 (36) and *Klebsiella pneumoniae* Kp08 (7). We use the first letter of the genus (E and K) to refer to these species throughout. The recipient strain is derived from the same isogenic strain as the *E. coli* donor strain but encodes additional chromosomal streptomycin resistance, providing a unique selectable marker to distinguish the donor and recipient hosts in both the cross-(K to E) and within-species (E to E) mating assays. The focal conjugative plasmid was used previously (37): plasmid F’42 from the IncF incompatibility group. A tetracycline resistance gene was cloned into the F’42 plasmid (38) and used as the selectable marker to distinguish plasmid-containing from plasmid-free hosts. This derived plasmid is referred to as ‘pF’ throughout.

### Conjugation Assays

Strains were inoculated into LB medium from frozen isogenic glycerol stocks and grown for approximately 24 hours. The plasmid-containing cultures were supplemented with 15 μg ml^-1^ tetracycline to maintain the plasmid. The saturated cultures were diluted 100-fold into LB medium to initiate another 24 hours of growth (to acclimate the previously frozen strains to laboratory conditions). The acclimated cultures were then diluted 10,000-fold into LB medium and incubated for strain specific times to ensure the cultures entered exponential growth (SI section 6b). The exponentially growing cultures were diluted by a factor specific to the donor-recipient pair (SI section 6e), mixed at equal volumes, and dispensed into 84 wells of a deep-well microtiter plate at 100 μl per well (Figure 5a black-bordered wells, these wells were the co-cultures used to estimate 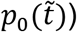. In an additional 3 wells, 130 μl (per well) of the mixture was dispensed and immediately 30 μl was removed to determine the initial densities (*D*_0_ and *R*_0_) via selective plating (Figure 5a black-bordered wells in top row). An additional 3 wells contained monocultures of the three strains. Specifically, 100 μl of donor, recipient and transconjugant cultures were placed in their own well (Figure 5a red-, blue- and purple-bordered wells, respectively, in the top left). Later in the assay, these monocultures determined if the transconjugant-selecting medium prohibited growth of both donors and recipients, while permitting growth of transconjugants. An additional 4 wells contained diluted monocultures of donors and recipients (2 wells each at 100 μl, Figure 5a red- and blue-bordered wells, respectively, in the top middle). These monocultures were used to create co-cultures (in empty wells, Figure 5a dash-bordered wells) during the assay itself (see below). The deep-well plate was incubated for a pre-determined time 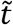 (SI section 6e), after which three events occurred in rapid succession. First, 30 μl was removed from each of the wells used to determine initial densities, to uncover the final densities (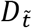 and 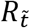) via selective plating (Figure 5b). Second, donor and recipient monocultures were mixed at equal volumes into the two empty wells (Figure 5b, gray arrows). At a later point in the assay, these two wells verified that new transconjugants did not form via conjugation after transconjugant-selecting medium was added. Third, 900 μl of transconjugant-selecting medium (7.5 μg ml^-1^ tetracycline and 25 μg ml^-1^ streptomycin; see SI section 6c and 6d) was added to all co-cultures used to estimate 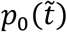 as well as relevant control wells (Figure 5c, yellow background). This medium disrupted new conjugation events—immediately by diluting cells then by inhibiting donors and recipients—while simultaneously selecting for transconjugant growth. The deep-well plate was incubated for 4 days, and the state of all wells (turbid or non-turbid) was recorded. For both mating assays in this study (i.e., cross- and within-species), this conjugation protocol was repeated 6 times.

For the cross-species mating, the SIM method was executed alongside the LDM method described above. The SIM approach was conducted for two incubation periods: a standard 24 hours and a truncated 5 hours. In an additional deep-well plate, 100 μl of the donor-recipient co-culture was dispersed into six wells, split into two groups of three wells each where each group corresponded to a different incubation period. To derive the SIM estimate for each incubation group, 30 μl was removed from each of the three wells in the group at the time point (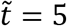 and 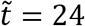) to determine the final donor 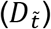, recipient 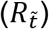, and transconjugant 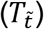 densities via selective plating. This protocol was repeated six times alongside the LDM replicates. Note the initial densities came from the matching LDM replicate. Similar to the LDM protocol, we ran a control to confirm that conjugation did not occur after co-cultures were exposed to transconjugant-selecting medium, but in this case, it was for agar plates instead of liquid medium. Specifically, for the first SIM replicate, an additional six donor monocultures and six recipient monocultures were initiated as above, again each split into two groups of three wells each. At each time point (5 and 24 hours), three new donor-recipient co-cultures were created in empty wells and, immediately plated on transconjugant-selecting agar at dilutions used to determine transconjugant densities. For this case, no transconjugant colonies formed (indicating that conjugation does not occur on the selective agar plate). We emphasize that this is a necessary step for any new system as post-plating conjugation has been reported (7, 13, 31).

For both the LDM and SIM approaches, the working assumption is that a cell will successfully establish a lineage under the appropriate selective conditions. As one example, a well with a single transconjugant will become turbid after incubation with transconjugant-selective medium. As another example, a donor cell on a donor-selecting agar plate will form a visible colony after incubation. A recent paper (39) has clearly demonstrated that this working assumption needs to be checked. In SI section 6, we offer adjustments to the protocols to improve the chances that this assumption holds. Additionally, we present ways to correct estimates if the assumption does not hold. In Figure 6, we used these corrections (see SI section 6 and 7 for details).

### Stochastics simulations

We used the Gillespie algorithm available in the GillesPy2 open-source Python package for stochastic simulations (40). We specified starting cell densities and parameters and simulated population dynamics using equations [1]-[3] for a set incubation time in a 1 ml culture volume. For each parameter setting, we simulated 10,000 populations and calculated the conjugation rate using the LDM and SIM estimates. Each estimate has different requirements for calculating the conjugation rate (Figure 3). The LDM estimate needs multiple populations to calculate 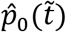; therefore, we reserved 100 populations to compute 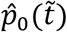 then one random population was used to calculate the initial and final cell densities. In other words, the 10,000 populations yielded 100 LDM estimates. In contrast, one simulated population yields one SIM estimate.

For the incubation time sweeps (Figure 3), the conjugation rate was estimated at 30-minute intervals up until the total population size reached 10^9^ cfu ml^-1^. A 30-minute interval was analyzed if at least 90 percent of the estimates were finite and non-zero. Notably, the 30-minute intervals occur over an earlier time range for the LDM estimate then for the SIM estimate due to the different estimate requirements. Given that these simulations are incubated until high population density is reached, the computational time for the Gillespie algorithm can be considerable. Therefore, 100 out of the 10,000 populations were incubated for the full incubation time (required to reach the saturated density of 10^9^ cfu ml^-1^) to provide SIM estimates over the time frame of interest. The remaining simulations were incubated for a truncated time frame until on average 100 transconjugants were generated to provide the populations needed to compute the 100 LDM estimates.

To compare across various parameter settings (Figure 4), a single incubation time was chosen. For each parameter setting, the incubation time 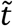 for the LDM estimate is set to the average *t*^*^. In addition, the incubation time for the SIM estimate is given by the time point for which an average of 50 transconjugants is reached. This choice resulted in a truncated SIM approach (i.e., 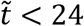). However, any estimate bias from a truncated simulation would be conservative relative to the standard SIM approach. At each incubation time, 10,000 simulated populations were used to calculate the estimate distribution.

## Data and Code Availability

All generated data and custom software are deposited in a GitHub repository (https://github.com/livkosterlitz/LDM).

## Acknowledgements

This work is supported by the National Institute of Allergy and Infectious Diseases Extramural Activities grant no. R01 AI084918 of the National Institutes of Health. O.K. is supported by the NSF Graduate Research Fellowship grant no. DGE-1762114. C.E. is supported by the NSF Graduate Research Fellowship. We thank Hannah Jordt and members of the Kerr and Top laboratories for useful suggestions on the manuscript.

## References

1. C. M. Thomas, K. M. Nielsen, Mechanisms of, and barriers to, horizontal gene transfer between bacteria. Nat. Rev. Microbiol.3, 711–721 (2005).

2. F. de la Cruz, J. Davies, Horizontal gene transfer and the origin of species: lessons from bacteria. Trends Microbiol.8, 128–133 (2000).

3. A. Norman, L. H. Hansen, S. J. Sørensen, Conjugative plasmids: vessels of the communal gene pool. Philos. Trans. R. Soc. Lond. B Biol. Sci.364, 2275–2289 (2009).

4. P. Mazodier, J. Davies, Gene transfer between distantly related bacteria. Annu. Rev. Genet.25, 147–171 (1991).

5. S. Redondo-Salvo, et al., Pathways for horizontal gene transfer in bacteria revealed by a global map of their plasmids. Nat. Commun.11, 3602 (2020).

6. A. K. Olesen, et al., IncHI1A plasmids potentially facilitate a horizontal flow of antibiotic resistance genes to pathogens in microbial communities of urban residential sewage. Mol. Ecol. (2022) https://doi.org/10.1111/mec.16346.

7. H. Jordt, et al., Coevolution of host-plasmid pairs facilitates the emergence of novel multidrug resistance. Nat Ecol Evol 4, 863–869 (2020).

8. J. Davies, Inactivation of antibiotics and the dissemination of resistance genes. Science 264, 375–382 (1994).

9. C. J. L. Murray, et al., Global burden of bacterial antimicrobial resistance in 2019: a systematic analysis. Lancet (2022) https://doi.org/10.1016/S0140-6736(21)02724-0.

10. B. R. Levin, F. M. Stewart, V. A. Rice, The kinetics of conjugative plasmid transmission: fit of a simple mass action model. Plasmid 2, 247–260 (1979).

11. L. Simonsen, D. M. Gordon, F. M. Stewart, B. R. Levin, Estimating the rate of plasmid transfer: an end-point method. J. Gen. Microbiol.136, 2319–2325 (1990).

12. A. J. Lopatkin, et al., Antibiotics as a selective driver for conjugation dynamics. Nat Microbiol 1, 16044 (2016).

13. J. H. Bethke, et al., Environmental and genetic determinants of plasmid mobility in pathogenic Escherichia coli. Sci Adv 6, eaax3173 (2020).

14. J. S. Huisman, et al., Estimating plasmid conjugation rates: a new computational tool and a critical comparison of methods. bioRxiv, 2020.03.09.980862 (2021).

15. T. Dimitriu, L. Marchant, A. Buckling, B. Raymond, Bacteria from natural populations transfer plasmids mostly towards their kin. Proc. Biol. Sci.286, 20191110 (2019).

16. A. San Millan, Evolution of Plasmid-Mediated Antibiotic Resistance in the Clinical Context. Trends Microbiol.26, 978–985 (2018).

17. L. Li, et al., Plasmids persist in a microbial community by providing fitness benefit to multiple phylotypes. ISME J.14, 1170–1181 (2020).

18. J. P. J. Hall, E. Harrison, D. A. Baltrus, Introduction: the secret lives of microbial mobile genetic elements. Philos. Trans. R. Soc. Lond. B Biol. Sci.377, 20200460 (2022).

19. S. E. Luria, M. Delbrück, Mutations of Bacteria from Virus Sensitivity to Virus Resistance. Genetics 28, 491–511 (1943).

20. A. R. Johnsen, N. Kroer, Effects of stress and other environmental factors on horizontal plasmid transfer assessed by direct quantification of discrete transfer events. FEMS Microbiol. Ecol.59, 718–728 (2007).

21. N. Fedoroff, W. Fontana, Genetic networks.Small numbers of big molecules.Science 297, 1129–1131 (2002).

22. W. Loftie-Eaton, et al., Compensatory mutations improve general permissiveness to antibiotic resistance plasmids. Nat Ecol Evol 1, 1354–1363 (2017).

23. J. B. Alderliesten, et al., Effect of donor-recipient relatedness on the plasmid conjugation frequency: a meta-analysis. BMC Microbiol.20, 135 (2020).

24. R. J. Sheppard, A. E. Beddis, T. G. Barraclough, The role of hosts, plasmids and environment in determining plasmid transfer rates: A meta-analysis. Plasmid 108, 102489 (2020).

25. C. Dahlberg, L. Chao, Amelioration of the cost of conjugative plasmid carriage in Eschericha coli K12. Genetics 165, 1641–1649 (2003).

26. W. Loftie-Eaton, et al., Contagious Antibiotic Resistance: Plasmid Transfer among Bacterial Residents of the Zebrafish Gut. Appl. Environ. Microbiol. 87(2021).

27. P. D. Lundquist, B. R. Levin, Transitory derepression and the maintenance of conjugative plasmids. Genetics 113, 483–497 (1986).

28. S. Bates, A. M. Cashmore, B. M. Wilkins, IncP plasmids are unusually effective in mediating conjugation of Escherichia coli and Saccharomyces cerevisiae: involvement of the tra2 mating system. J. Bacteriol.180, 6538–6543 (1998).

29. A. E. Dewar, et al., Plasmids do not consistently stabilize cooperation across bacteria but may promote broad pathogen host-range. Nat Ecol Evol 5, 1624–1636 (2021).

30. K. R. Philipsen, L. E. Christiansen, H. Hasman, H. Madsen, Modelling conjugation with stochastic differential equations. J. Theor. Biol.263, 134–142 (2010).

31. E. Smit, J. D. van Elsas, Determination of plasmid transfer frequency in soil: Consequences of bacterial mating on selective agar media. Curr. Microbiol.21, 151–157 (1990).

32. De Gelder Leen, Vandecasteele Frederik P. J., Brown Celeste J., Forney Larry J., Top Eva M., Plasmid Donor Affects Host Range of Promiscuous IncP-1β Plasmid pB10 in an Activated-Sludge Microbial Community. Appl. Environ. Microbiol.71, 5309–5317 (2005).

33. A. Kottara, J. P. J. Hall, M. A. Brockhurst, The proficiency of the original host species determines community-level plasmid dynamics. FEMS Microbiol. Ecol.97 (2021).

34. A. Kottara, L. Carrilero, E. Harrison, J. P. J. Hall, M. A. Brockhurst, The dilution effect limits plasmid horizontal transmission in multispecies bacterial communities. Microbiology 167(2021).

35. J. D. van Elsas, M. J. Bailey, The ecology of transfer of mobile genetic elements. FEMS Microbiol. Ecol.42, 187–197 (2002).

36. W. Loftie-Eaton, et al., Evolutionary Paths That Expand Plasmid Host-Range: Implications for Spread of Antibiotic Resistance. Mol. Biol. Evol.33, 885–897 (2016).

37. R. J. F. Haft, et al., General mutagenesis of F plasmid TraI reveals its role in conjugative regulation. J. Bacteriol.188, 6346–6353 (2006).

38. H. L. Jordt, “Of E. coli and classrooms: Stories of persistence.” (2019).

39. H. K. Alexander, R. C. MacLean, Stochastic bacterial population dynamics restrict the establishment of antibiotic resistance from single cells. Proc. Natl. Acad. Sci. U. S. A.117, 19455–19464 (2020).

40. J. H. Abel, B. Drawert, A. Hellander, L. R. Petzold, GillesPy: A Python Package for Stochastic Model Building and Simulation. IEEE Life Sci Lett 2, 35–38 (2016).

