## Supplemental Information for "Estimating the rate of plasmid transfer with an adapted Luria–Delbrück fluctuation analysis"

**This PDF file includes:**
Supplementary text
SI Figures 1 to 10
SI Tables 1 to 8

### SI section 1 : Overview of approaches to estimate the conjugation rate.

#### *SI section 1a : Overview of theoretical frameworks*

In this section, we highlight three key methods for estimating conjugation rate. While outlining the theoretical frameworks, we highlight the key distinctions and theoretical assumptions of each approach. Levin *et. al.* (1) introduced a simple mathematical model describing the change in density of donors, recipients and transconjugants over time (given by dynamic variables $D_t$, $R_t$, and $T_t$, respectively). In this model, each population type grows exponentially at the same growth rate $\psi$. In addition, the transconjugant density increases because of conjugation events both from donors to recipients and from existing transconjugants to recipients at the same conjugation rate $\gamma$. The recipient density decreases due to these conjugation events. The densities of these dynamic populations are described by the following differential equations (where the $t$ subscript is dropped from the dynamic variables for notational convenience):

$$\frac{dD}{dt} = \psi D, \qquad [1.1]$$

$$\frac{dR}{dt} = \psi R - \gamma DR - \gamma TR, \qquad [1.2]$$

$$\frac{dT}{dt} = \psi T + \gamma DR + \gamma TR. \qquad [1.3]$$

Equations [1.1]-[1.3] contains four notable assumptions. First, conjugation is described by mass-action kinetics, where conjugation events are proportional to the product of donor and recipient cell densities, which is a reasonable assumption in well-mixed liquid cultures (1). Second, the model assumes a negligible rate of plasmid loss, a process whereby a dividing plasmid-containing cell produces one plasmid-containing daughter cell and one plasmid-free daughter. These first two assumptions exist in all the conjugation rate estimates we discuss. Third, the growth rate is the same for all cell types (i.e., in the language of equations [1]-[3], $\psi_D = \psi_R = \psi_T = \psi$). Fourth, the plasmid conjugation rate is the same from donors to recipients as from transconjugants to recipients (i.e., in the language of equations [1]-[3], $\gamma_D = \gamma_T = \gamma$). More specifically, equations [1.1]-[1.3] are a special case of equations [1]-[3] where growth and conjugation is assumed to be homogeneous across strains.

Popular rate estimation methods solved the set of ordinary differential equations from the Levin *et. al.* model (or a variation) to find an estimate for the conjugation rate $\gamma$. The various methods differ by the assumptions used to find the analytical solution. Levin *et. al.* was the first to derive an estimate for the conjugation rate ($\gamma$) by making three additional simplifying assumptions. First, the change in cell density of donors due to growth is assumed to be negligible. Likewise, the change in cell density of recipients due to growth and to conjugation (i.e., transformation into transconjugants) is assumed to be negligible. Finally, transconjugants are assumed to be rare in the population resulting in negligible growth of transconjugants and insignificant conjugation from transconjugants to recipients. Thus, the increase in transconjugant cell density is through plasmid conjugation from donors to recipients ($2^{nd}$ term equation [1.3]). Using these simplifying assumptions, Levin *et. al.* solved for an expression of the conjugation rate in terms of the density of donors, recipients, and transconjugants ($D_{\tilde{t}}$, $R_{\tilde{t}}$, and $T_{\tilde{t}}$, respectively) after a period of incubation $\tilde{t}$ (see the GitHub Appendix I for a few different approaches to the derivation).

$$\gamma = \frac{T_{\tilde{t}}}{D_{\tilde{t}} R_{\tilde{t}} \tilde{t}}. \quad [1.4]$$

We label the expression in equation [1.4] as the "TDR" estimate for the conjugation rate, where the letters in this acronym come from the dynamic variables used in the estimate. Besides the model assumes homogenous growth rates and conjugation rates, the most notable assumption used in the TDR derivation is that there is little to no change in the population densities due to growth. Thus, laboratory implementation that respects this assumption can be difficult (see SI section 1b for details). Regardless, TDR is a commonly used estimate (1–4).

Simonsen *et. al.* derived the other most widely used estimate for conjugation rate $\gamma$, which importantly expands application beyond the TDR method by allowing for population growth (5). Indeed, they allowed for the rate of population growth to change with the level of a resource in the environment, adding a dynamic variable for the resource concentration. In addition, the conjugation rate can also change with the resource concentration. The authors focus on a case where both growth and conjugation rates vary with resource concentration according to the Monod function. This choice was informed by experimental results showing that cells enter stationary phase and conjugation ramps down to a negligible level as resources are depleted (1). This pattern occurs for various plasmid incompatibility groups, but not all (6). Simonsen *et. al.* used this updated model to derive an estimate for plasmid conjugation rate (see GitHub Appendix II for the derivation).

$$\gamma_D = \psi \ln\left(1 + \frac{T_{\tilde{t}}}{R_{\tilde{t}}} \frac{N_{\tilde{t}}}{D_{\tilde{t}}}\right) \frac{1}{(N_{\tilde{t}} - N_0)}. \quad [1.5]$$

Equation [1.5] we refer to as the "SIM" estimate for conjugation rate throughout the manuscript, where SIM stands for "Simonsen *et. al.* Identicality Method" since the underlying model assumes that all strains are *identical* with regards to growth, and donors and transconjugants are *identical* with regards to conjugation rate. The SIM estimate involves measuring the density of the initial population ($N_0$), and the final density of donors ($D_{\tilde{t}}$), recipients ($R_{\tilde{t}}$), transconjugants ($T_{\tilde{t}}$), and the total population ($N_{\tilde{t}}$) after the incubation time $\tilde{t}$. The SIM estimate is a commonly used since it allows for the use of batch culture in the laboratory (see SI section 1b for details). Thus, it circumvents the constraints of the laboratory implementation of TDR; however, the underlying model holds the same assumptions as before: homogeneous growth rates and conjugation rates.

Huisman *et. al.* recently updated the SIM model, further extending its application by relaxing the assumption of identical growth and transfer rates for all strains (7). Specifically, the authors introduced population specific growth rates for donors, recipients, and transconjugants ($\psi_D$, $\psi_R$, and $\psi_T$, respectively) and population specific conjugation rates for donors and transconjugants ($\gamma_D$ and $\gamma_T$). Huisman *et. al.* made three additional simplifying assumptions. First, conjugation and growth rates are assumed to be constant until resources are depleted, eliminating the additional resource concentration equation added in the SIM approach. Second, the increase in recipients due to growth greatly outpaces the decrease in recipients due to conjugation (i.e., $\psi_R R \gg \gamma_D DR + \gamma_T TR$). Third, the increase in transconjugants due to growth or plasmid conjugation from donors to recipients greatly outpaces the increase in transconjugants due to plasmid conjugation from transconjugants to recipients (i.e., $\psi_T T + \gamma_D DR \gg \gamma_T TR$). These model conditions are reasonable if the system starts with donors and recipients present but transconjugants are absent, the system is tracked for a short period of time $\tilde{t}$, conjugation rates are low relative to growth rates, and the transconjugant conjugation rate ($\gamma_T$) is not much higher than the donor conjugation rate ($\gamma_D$). With these added assumptions, equations [1.1]-[1.3] can be reformulated as the following *approximate* system of equations:

$$\frac{dD}{dt} = \psi_D D, \quad [1.6]$$

$$\frac{dR}{dt} = \psi_R R, \quad [1.7]$$

$$\frac{dT}{dt} = \psi_T T + \gamma_D D R, \quad [1.8]$$

Huisman *et. al.* used these equations to derive an estimate for the donor conjugation rate

$$\gamma_D = (\psi_D + \psi_R - \psi_T)\frac{T_{\tilde{t}}}{\left(D_{\tilde{t}} R_{\tilde{t}} - D_0 R_0 e^{\psi_T \tilde{t}}\right)}. \quad [1.9]$$

where different cell types now can have different growth rates (see GitHub Appendix III for the derivation). We term equation [1.9] as the ASM estimate for donor conjugation rate, where ASM stands for “Approximate Simonsen *et. al.* Method”.

For all methods (TDR, SIM, ASM, and LDM), we summarize model variables and parameters in SI Table 1. In addition, all variables used in the conjugation estimates are in SI Table 2. Lastly, all assumptions underlying each estimate are in SI Table 3.

**SI Table 1: Variables and parameters used in plasmid dynamic models.**

| Variable/ Parameter | Description | Relevant Estimate(s) | Units |
|---|---|---|---|
| $D$ | Donor density | TDR, SIM, ASM, LDM | $\frac{\text{cfu}}{\text{ml}}$ |
| $R$ | Recipient density | TDR, SIM, ASM, LDM | |
| $T$ | Transconjugant density | TDR, SIM, ASM, LDM | |
| $\psi$ | Growth rate (not population specific) | TDR, SIM | $\frac{1}{\text{hr}}$ |
| $\psi_D$ | Donor growth rate | ASM, LDM | |
| $\psi_R$ | Recipient growth rate | ASM, LDM | |
| $\psi_T$ | Transconjugant growth rate | ASM, LDM | |
| $\gamma$ | Conjugation rate (not population specific) | TDR, SIM | $\frac{\text{ml}}{\text{cfu} \cdot \text{hr}}$ |
| $\gamma_D$ | Donor-recipient conjugation rate | ASM, LDM | |
| $\gamma_T$ | Transconjugant-recipient conjugation rate | ASM, LDM | |

**SI Table 2: Variables and parameters used to estimate* conjugation rate**

| Variable/Parameter | Description | Relevant Estimate | Units |
|---|---|---|---|
| $\tilde{t}$ | Incubation time (final sampling time) | TDR, SIM**, ASM, LDM | hr |

| | | | |
|---|---|---|---|
| $D_0, R_0$ | Initial donor and recipient densities | ASM, LDM | $\frac{\text{cfu}}{\text{ml}}$ |
| $D_{\tilde{t}}, R_{\tilde{t}}$ | Final donor and recipient densities | TDR, SIM, ASM, LDM | |
| $T_{\tilde{t}}$ | Final transconjugant density | TDR, SIM, ASM | |
| $N_0, N_{\tilde{t}}$ | Initial and final total population density | SIM | |
| $\psi_T$ | Transconjugant growth rate | ASM | $\text{hr}^{-1}$ |
| $p_0(\tilde{t})$ | Probability a population has no transconjugants | LDM | |
| * The laboratory estimates are used here (see SI section 1b)<br>** If the SIM assay is conducted on exponentially growing cultures (see SI section 1c) $\tilde{t}$ along with $N_0$ and $N_{\tilde{t}}$ can be used to estimate $\psi$ (otherwise, an independent estimate of $\psi$ is needed). | | | |

**SI Table 3: Summary of modeling assumptions.**

| Assumption | TDR | SIM | ASM | LDM |
|---|---|---|---|---|
| Conjugation events follow mass-action kinetics | X | X | X | X |
| The plasmid loss rate of the focal plasmid is zero | X | X | X | X |
| The cell populations do not change in size due to growth | X | | | |
| Processes of conjugation and growth are not resource dependent* | X | | X | X |
| The cell populations grow exponentially (i.e., constant growth rate) | | | X | X |
| The growth rate is identical for all cell types | X | X | | |
| The transconjugant conjugation rate is not high relative to the donor conjugation rate | X | X | X | |
| * The SIM model can incorporate resource-dependent growth and conjugation because (1) growth and transfer rates are homogeneous and (2) the functional form for resource dependence is the same for growth and transfer. | | | | |

*SI section 1b : Alternative laboratory forms for conjugation estimates*

Often the conjugation estimates can be re-written into a form of the equation which is more amenable to laboratory implementation. Here we walk through rearranging the equations for a subset of the estimates.

For the SIM estimate, we start with equation [1.5]. If the entire period from $t = 0$ to $t = \tilde{t}$ involves exponential growth, then $N_{\tilde{t}} = N_0 e^{\psi \tilde{t}}$. In such a case, $\psi = (1/\tilde{t})\ln(N_{\tilde{t}}/N_0)$. We arrive at the alternative laboratory form for SIM

$$\gamma = \frac{1}{\tilde{t}}\left[\ln\left(1 + \frac{T_{\tilde{t}}}{R_{\tilde{t}}}\frac{N_{\tilde{t}}}{D_{\tilde{t}}}\right)\right]\frac{\ln N_{\tilde{t}} - \ln N_0}{N_{\tilde{t}} - N_0}. \quad [1.10]$$

We note that equation [1.9] is appropriate for some "truncated" versions of the SIM approach, but not generally applicable to the standard overnight version in which the culture does not grow exponentially across the entire assay.

To rearrange the ASM estimate, we start with equation [1.9]. While the equations $\psi_D = (1/\tilde{t})\ln(D_{\tilde{t}}/D_0)$ and $\psi_R = (1/\tilde{t})\ln(R_{\tilde{t}}/R_0)$ again provide estimates on donor and recipient growth rates, we cannot express the transconjugant growth rate ($\psi_T$) as a simple expression of time and initial/final densities of members of the mating culture. However, data from a transconjugant monoculture supply an estimate for this parameter. Thus, we arrive at the laboratory form for ASM

$$\gamma_D = \left\{\frac{1}{\tilde{t}}(\ln D_{\tilde{t}}R_{\tilde{t}} - \ln D_0R_0) - \psi_T\right\}\frac{T_{\tilde{t}}}{\left(D_{\tilde{t}}R_{\tilde{t}} - D_0R_0e^{\psi_T\tilde{t}}\right)}. \qquad [1.11]$$

To rearrange the LDM estimate, we start with equation [11]. Since $D_{\tilde{t}} = D_0e^{\psi_D\tilde{t}}$ and $R_{\tilde{t}} = R_0e^{\psi_R\tilde{t}}$, it is the case that $\psi_D = (1/\tilde{t})\ln(D_{\tilde{t}}/D_0)$ and $\psi_R = (1/\tilde{t})\ln(R_{\tilde{t}}/R_0)$. So, we have

$$\gamma_D = \frac{1}{\tilde{t}}\ln p_0(\tilde{t})\frac{\ln\left(\frac{D_{\tilde{t}}}{D_0}\right) + \ln\left(\frac{R_{\tilde{t}}}{R_0}\right)}{D_0R_0 - D_{\tilde{t}}R_{\tilde{t}}}$$

After rearrangement, we have

$$\gamma_D = \frac{1}{\tilde{t}}\{-\ln p_0(\tilde{t})\}\frac{\ln(D_{\tilde{t}}R_{\tilde{t}}) - \ln(D_0R_0)}{D_{\tilde{t}}R_{\tilde{t}} - D_0R_0}.$$

In the laboratory, we measure an estimate ($\hat{p}_0(\tilde{t})$) of the probability that a population has no transconjugants ($p_0(\tilde{t})$) which is simply the fraction of the populations (i.e., parallel cultures) that have no transconjugants at the incubation time $\tilde{t}$. In addition, if a 1 ml volume is not used for each mating culture (assuming that all cell densities are measured in $\mathrm{cfu/ml}$ units), then we must add a correction factor $f$ (see SI section 5 for details and an example). Thus, we arrive at the laboratory form for the LDM, which is equation [13].

$$\gamma_D = \frac{f}{\tilde{t}}[-\ln\hat{p}_0(\tilde{t})]\frac{\ln D_{\tilde{t}}R_{\tilde{t}} - \ln D_0R_0}{D_{\tilde{t}}R_{\tilde{t}} - D_0R_0}$$

*SI section 1c : Overview of laboratory implementations*

In this section, we compare the laboratory implementations of the various estimates: TDR, SIM, and ASM. Each method is explained either as recommended by its authors or the most simplified protocol to acquire the information for the estimate. For each, we describe proper laboratory implementation for the approaches based on the model and derivation assumptions used to acquire the estimate. Note in this section, we do not explore the assumptions that are violated due to the biological entities being tested (i.e., specific species or plasmids) which can result in violations such as unequal conjugation rates or growth rates. These are explored in the main text and SI section 4 via stochastic simulations. Thus, we focus solely on the parameters under the experimenter's control. For ease of reference, key implementation differences are highlighted in SI Table 4.

The TDR estimate has a simple form (equation [1.4]). Donors and recipients are mixed in non-selective growth medium and incubated for a specified time $\tilde{t}$. Typically, densities after the incubation time are determined using selective plating. The derivation assumes the density in donors and recipients does not change due to growth which sets specific constraints on the implementation of this approach. In the original study, Levin *et. al.*, used a chemostat to keep the population constant (1). Other studies shorten the incubation time $\tilde{t}$ such that population growth is negligible and use various laboratory tools to detect the small number of transconjugants (3, 4).

The SIM estimate is not built on an assumption of unchanging population densities. Donor and recipient populations in exponential phase are mixed in non-selective growth medium. The initial population density ($N_0$) is determined by dilution plating on non-selective medium. After the mating mixture is incubated (for a period of $\tilde{t}$), the final densities ($D_{\tilde{t}}$, $R_{\tilde{t}}$, $T_{\tilde{t}}$, and $N_{\tilde{t}}$) are determined by dilution plating on selective and non-selective media. To implement SIM as written in equation [1.10] (see SI section 1b), the specified incubation time $\tilde{t}$ must occur well before stationary phase is reached to collect proper data for estimating the population growth rate ($\psi = (\ln N_{\tilde{t}} - \ln N_0)/\tilde{t}$). There is an alternative option for implementing the SIM using equation [1.5]. The donor and recipient populations are mixed and incubated under batch culture conditions (specifically exponential and stationary phase). However, the (maximum) population growth rate ($\psi$) needs to be determined with two additional samplings from the mixed population at times $t_a$ and $t_b$, both occurring *within exponential phase:*

$$\psi = \frac{\ln (N_{t_b}/N_{t_a})}{t_b - t_a} \qquad [1.12]$$

The population densities $N_{t_a}$ and $N_{t_b}$ can be estimated either through colony counts from plating or optical density from a spectrophotometer. Either way, the timing of exponential phase is important for this approach and at least some analysis during this phase is required regardless of the implementation strategy.

For the ASM estimate, donor and recipient populations in exponential phase are mixed in non-selective medium. Initial densities ($D_0$ and $R_0$) are determined by plating dilutions on the appropriate selective media. After the donor-recipient co-culture incubates for a specified time ($\tilde{t}$), final densities ($D_{\tilde{t}}$, $R_{\tilde{t}}$, and $T_{\tilde{t}}$) are determined by plating dilutions on the appropriate selective media. From the transconjugant-selecting agar plates, a transconjugant clone is incubated in monoculture then sampled twice (at times $t_a$ and $t_b$) in exponential phase to measure the transconjugant growth rate ($\psi_T = \ln(T_{t_b}/T_{t_a})/(t_b - t_a)$). The authors point out a critical consideration for proper implementation of the ASM is the incubation time $\tilde{t}$. Not only is sampling in exponential phase important, but if the incubation time $\tilde{t}$ is too long and passes a critical time ($t_{crit}$) the approximations used to derive the ASM break down. To avoid overshooting $t_{crit}$, the authors recommend sampling as soon as measurable transconjugants arise. To determine that the incubation time used was below the critical time ($t_{crit}$), a second assay is recommended by the authors to measure the transconjugant conjugation rate $\gamma_T$, which will determine if the original incubation time $\tilde{t}$ was below $t_{crit}$ for measuring the donor conjugation rate. This second assay would have the transconjugant clone become the donor in the mixture, while a newly marked recipient must be used so that donors and recipients can be distinguished using selective plating.

Each method has aspects of implementation in common. Each one shares the basic approach of mixing donors and recipients over some incubation time $\tilde{t}$. Each estimate requires reliable selectable markers to differentiate donors, recipients, and transconjugants. However, all estimates have some constraints on initial densities and time of measurement. This can occur because the experimenter needs to capture conjugation events (all estimates require this), avoid population growth (TDR), or keep growth exponential (ASM, and at least parts of SIM). Even so, each method has clear distinctions. The most notable is the incubation time $\tilde{t}$ (i.e., the end of the assay). The TDR method is constrained to conditions where no change in population size due to growth can occur. For SIM, initial and final sampling are not constrained to a particular phase of growth; however, measurement of the growth rate must occur during the exponential growth phase. For ASM, initial sampling is in early exponential phase, and the final sampling needs to occur during a specific time window. In other words, the assay needs to be long enough that measurable transconjugants appear, but short enough so that assumptions are not violated (which can occur if transconjugant density becomes too large).

**SI Table 4: Comparison of implementations.**

| Summary | TDR | SIM | ASM | LDM |
|---|---|---|---|---|
| Assay conditions minimizing the change in density due to growth | X | | | |
| Minimize incubation time necessary for producing transconjugants | | | X | |
| An incubation time results in a subset of parallel populations having no transconjugants | | | | X |
| Assay occurs over a period of exponential cell growth | | X* | X | X |
| Assay requires multiple parallel mating cultures per replicate | | | | X |
| Assay requires a measurement of transconjugant density | X | X | X | |
| Assay requires a measurement of population growth rate | | X* | | |
| Assay requires a measurement of transconjugant growth rate | | | X | |
| * For the SIM assay, either the entire assay is conducted over exponentially growing cultures or an independent estimate for (maximum) population growth rate is needed. | | | | |

### SI section 2 : Derivation of $p_0(\tilde{t})$ for the LDM estimate

In this section, we will continue to assume an experimental volume of 1 ml for the co-culture such that the density of cells per ml is equivalent to the cell count numerically. We will not explicitly track units in this section, but we deal with the case of an arbitrary experimental volume in SI section 5.

We define $p_n(t)$ to be the probability that there are $n$ transconjugants at time $t$, where $n$ is a non-negative integer (i.e., $p_n(t) = \Pr\{T_t = n\}$). We focus here on the probability that transconjugants are absent (namely, where $n = 0$) and derive an expression for $p_0(t)$. By definition $p_0(t + \Delta t) = \Pr\{T_{t+\Delta t} = 0\}$. However, $T_{t+\Delta t} = 0$ implies $T_t = 0$, so we can write

$$p_0(t + \Delta t) = \Pr\{(T_{t+\Delta t} = 0) \cap (T_t = 0)\} = \Pr\{T_{t+\Delta t} = 0 \mid T_t = 0\} \Pr\{T_t = 0\},$$

Given that $p_0(t) = \Pr\{T_t = 0\}$, we can use equation [9] to write the following time-increment recursion for $p_0(t)$:

$$p_0(t + \Delta t) = (1 - \gamma_D D_t R_t \Delta t) p_0(t).$$

This can be simplified as follows

$$\frac{p_0(t + \Delta t) - p_0(t)}{\Delta t} = -\gamma_D D_t R_t p_0(t).$$

Taking the limit as $\Delta t \to 0$ gives

$$\lim_{\Delta t \to 0} \frac{p_0(t + \Delta t) - p_0(t)}{\Delta t} = \frac{dp_0(t)}{dt}.$$

Therefore, we have the following differential equation:

$$\frac{dp_0(t)}{dt} = -\gamma_D D_t R_t p_0(t).$$

We are assuming $D_t = D_0 e^{\psi_D t}$ and $R_t = R_0 e^{\psi_R t}$. We note that these assumptions are reasonable if the densities of donors and recipients are reasonably large and the rate of transconjugant generation per recipient cell ($\gamma_D D_t + \gamma_T T_t$, or if $T_t = 0$, simply $\gamma_D D_t$) remains very small relative to per capita recipient growth rate ($\psi_R$). Under these assumptions, our differential equation becomes:

$$\frac{dp_0(t)}{dt} = -\gamma_D D_0 R_0 e^{(\psi_D+\psi_R)t} p_0(t).$$

We solve this differential equation via separation of variables, integrating from 0 to our incubation time of interest $\tilde{t}$:

$$\int_0^{\tilde{t}} \frac{dp_0(t)}{p_0(t)} = \int_0^{\tilde{t}} -\gamma_D D_0 R_0 e^{(\psi_D+\psi_R)t} dt,$$

$$\ln p_0(t)\big|_0^{\tilde{t}} = \frac{-\gamma_D D_0 R_0}{\psi_D + \psi_R} e^{(\psi_D+\psi_R)t}\Big|_0^{\tilde{t}},$$

$$\ln p_0(\tilde{t}) - \ln p_0(0) = \frac{-\gamma_D D_0 R_0}{\psi_D + \psi_R} e^{(\psi_D+\psi_R)\tilde{t}} - \frac{-\gamma_D D_0 R_0}{\psi_D + \psi_R}.$$

Given that $p_0(0) = 1$,

$$\ln p_0(\tilde{t}) = \frac{-\gamma_D D_0 R_0}{\psi_D + \psi_R}\left(e^{(\psi_D+\psi_R)\tilde{t}} - 1\right),$$

$$p_0(\tilde{t}) = \exp\left\{\frac{-\gamma_D D_0 R_0}{\psi_D + \psi_R}\left(e^{(\psi_D+\psi_R)\tilde{t}} - 1\right)\right\},$$

which is equation [10].

### SI section 3 : Derivation of mutation rate from the Luria-Delbrück experiment

Here we derive the classic estimate of mutation rate from Luria and Delbrück. We assume that there is a population of wild-type cells that grow according to the following equation:

$$N_t = N_0 e^{\psi_N t}, \quad [3.1]$$

where $N_t$ is the number of wild type cells at time $t$ and $\psi_N$ is the per capita growth rate. The wild-type population dynamics are assumed to be deterministic (a reasonable assumption if the initial population size is reasonably large, i.e., $N_0 \gg 0$). We are also ignoring the loss of wild-type cells to mutational transformation, but this omission is reasonable if the mutation rate is very small relative to per capita wild-type growth rate.

Let the number of mutants be given by a random variable $M_t$. This variable takes on non-negative integer values. For a very small interval of time, $\Delta t$, the current number of mutants will either increase by one or remain constant. The probabilities of each possibility are given as follows:

$$\Pr\{M_{t+\Delta t} = M_t + 1\} = \mu N_t \Delta t + \psi_M M_t \Delta t, \quad [3.2]$$

$$\Pr\{M_{t+\Delta t} = M_t\} = 1 - (\mu N_t + \psi_M M_t)\Delta t. \quad [3.3]$$

The two terms on the right-hand side of equation [3.2] give the ways that a mutant can be generated. The first term measures the probability that a wild-type cell undergoes a mutation ($\mu$ is the mutation rate). The second term gives the probability that a mutant cell divides and produces two mutant cells ($\psi_M$ is the mutant growth rate). Equation [3.3] is the probability that neither of these processes occur.

Analogous to the procedure in SI section 2 (with $p_0(t) = \Pr\{M_t = 0\}$):

$$p_0(t + \Delta t) = (1 - \mu N_t \Delta t)p_0(t).$$

By rearranging, taking the limit as $\Delta t \to 0$, and utilizing equation [3.1], we have

$$\frac{dp_0(t)}{dt} = -\mu N_0 e^{\psi_N t} p_0(t).$$

This differential equation can be solved in an analogous way as well

$$\int_0^{\tilde{t}} \frac{dp_0(t)}{p_0(t)} = \int_0^{\tilde{t}} -\mu N_0 e^{\psi_N t} dt,$$

$$\ln p_0(t)|_0^{\tilde{t}} = \frac{-\mu N_0}{\psi_N} e^{\psi_N t}\Big|_0^{\tilde{t}},$$

$$\ln p_0(\tilde{t}) - \ln p_0(0) = \frac{-\mu N_0}{\psi_N} e^{\psi_N \tilde{t}} - \frac{-\mu N_0}{\psi_N}.$$

Because we assume $M_0 = 0$, we must have $p_0(0) = 1$, and

$$\ln p_0(\tilde{t}) = \frac{-\mu N_0}{\psi_N}\left(e^{\psi_N \tilde{t}} - 1\right).$$

Solving for the mutation rate $\mu$

$$\mu = -\ln p_0(\tilde{t}) \frac{\psi_N}{N_0\left(e^{\psi_N \tilde{t}} - 1\right)},$$

which is equation [12]. This equation can also be expressed as:

$$\mu = -\ln p_0(\tilde{t}) \frac{\psi_N}{N_{\tilde{t}} - N_0}.$$

To recover Luria and Delbrück's original formulation, consider a new time variable $z$ defined as follows:

$$z = \frac{t}{t_d/\ln 2},$$

where $t_d$ is the period required for population doubling during exponential growth. Because

$$N_0 e^{\psi_N t} = N_0 2^{t/t_d},$$

it is the case that

$$\psi_N = \frac{1}{t_d/\ln 2}.$$

Therefore, $z = \psi_N t$ and the equation $N_t = N_0 e^{\psi_N t}$ can be expressed as

$$N_z = N_0 e^z$$

Performing the same analysis on this new equation gives the original formulation (their equations [4] and [5], where our $\mu$ is given by their "a" and our $\tilde{z}$ is given by their "t"):

$$\mu = \frac{-\ln p_0(\tilde{z})}{N_{\tilde{z}} - N_0}.$$

### SI section 4 : Extended Simulation Results

#### *SI section 4a : Extended stochastic simulation methods*

To systematically explore the effects of heterogeneous growth and conjugation rates (as well as non-zero rates of plasmid loss) on the accuracy and precision of estimating the donor conjugation rate ($\gamma_D$), we developed a stochastic simulation framework using the Gillespie algorithm. We ran sets of simulations sweeping through parameter values. Each simulation examined a biological process (i.e., growth, conjugation) in isolation by manipulating one or two of the relevant parameters. For SI section 4b-d, we used a "baseline" set of parameters ($\psi_D = \psi_R = \psi_T = 1$, and $\gamma_D = \gamma_T = 1 \times 10^{-6}$) and initial densities ($D_0 = R_0 = 1 \times 10^2$ and $T_0 = 0$) unless otherwise indicated. For each initial parameter setting, we simulated 10,000 parallel populations and calculated the conjugation rate using each method (TDR, SIM, ASM, and LDM). The incubation time selection criteria used for the SIM estimate was also used for the TDR and ASM estimates (see Materials and Methods). However, given that all our simulated populations *increase in size* over the incubation time, a fundamental assumption of the TDR approach is broken for all the runs (i.e., no change in the density due to growth). The TDR estimate was included to be comprehensive (and illustrate that violation of the no growth assumption leads to systemic bias). Given that the Gillespie algorithm is computationally expensive and the large number of simulations needed to sweep through parameters, we chose low initial densities and a high conjugation rate for the baseline condition. In SI section 4e, we demonstrate that the trends shown for the baseline condition are also observed with more realistic parameter values and higher initial densities.

*SI section 4b : The effect of unequal growth rates*

We expanded the analysis used in Figure 4a by calculating a conjugation rate estimate with two additional estimates, TDR and ASM (SI Figure 1a). We simulated an additional biological scenario (SI Figure 1b) in which growth rates differ due to the host where recipients and transconjugants grow faster ($\psi_D < \psi_T = \psi_R$) or slower ($\psi_D > \psi_T = \psi_R$) than the donors. This captures the situation in which the recipient (and therefore transconjugant) is a different strain or species from the donor and differs in growth rate. Like the conclusions drawn from Figure 4 with the effects of plasmid carriage, the LDM exhibited high accuracy and precision relative to other metrics.

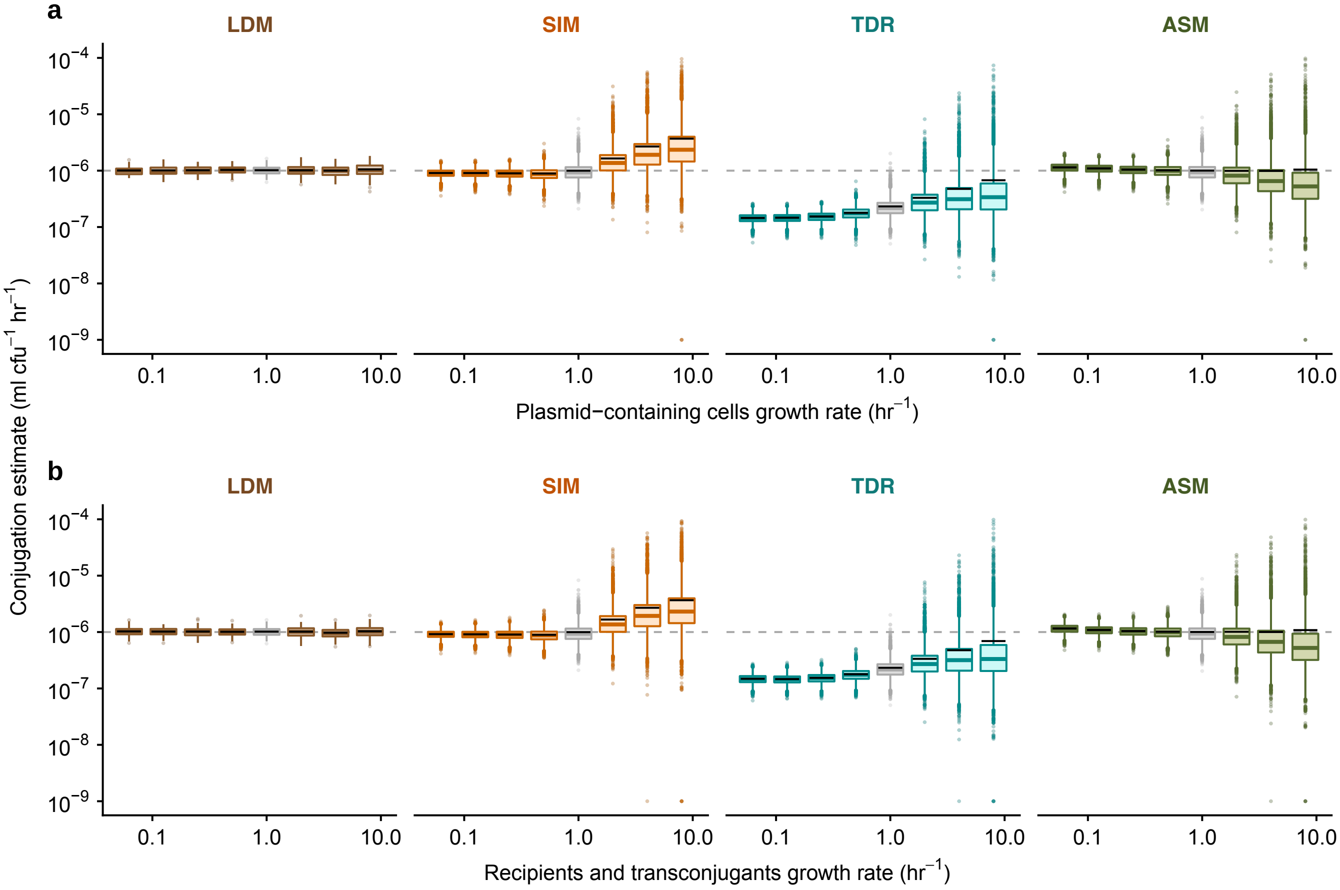


**SI Figure 1: The effect of heterogenous growth rates on estimating conjugation rate.** The Gillespie algorithm was used to simulate population dynamics. Donor conjugation rate for each parameter combination was estimated using 10,000 simulations (summarized using boxplots with the same graphical convention as in Figure 3). The gray dashed line indicates the true value for the donor conjugation rate (here, $10^{-6}$). The boxes in gray indicate the baseline parameter setting, and all colored boxes represent deviation of one or two parameters from baseline. The baseline parameter values were $\psi_D = \psi_R = \psi_T = 1$, and $\gamma_D = \gamma_T = 10^{-6}$. The dynamic variables were initialized with $D_0 = R_0 = 10^2$ and $T_0 = 0$. All incubation times are short but are specific to each parameter setting (see Materials and Methods for details). The LDM, SIM, TDR, and ASM estimates are in separate plots with estimate specific colors (brown, orange, cyan, and green, respectively). Zero estimates were set to $10^{-9}$ (the lowest y-value) for plotting on a log axis. (a) Unequal growth rates were explored over a range of growth rates for the plasmid-bearing strains, namely $\psi_D = \psi_T \in$ {0.0625, 0.125, 0.25, 0.5, 1, 2, 4, 8}. (b) Unequal growth rates were explored over a range of growth rates for the recipients and transconjugants, namely $\psi_R = \psi_T \in$ {0.0625, 0.125, 0.25, 0.5, 1, 2, 4, 8}.

In addition, we explored the effect of a single population (donor, recipient, and transconjugant) growing faster or slower in isolation (SI Figure 2). Notably, a faster transconjugant growth rate led to very large variance with the other metrics (TDR, SIM, and ASM). Therefore, some parameter settings shown in SI Figure 2c have a large proportion of the simulations yielding a zero estimate at the specific incubation time. Given the log-axis, the zero estimates were placed at the lowest y-value for plotting purposes.

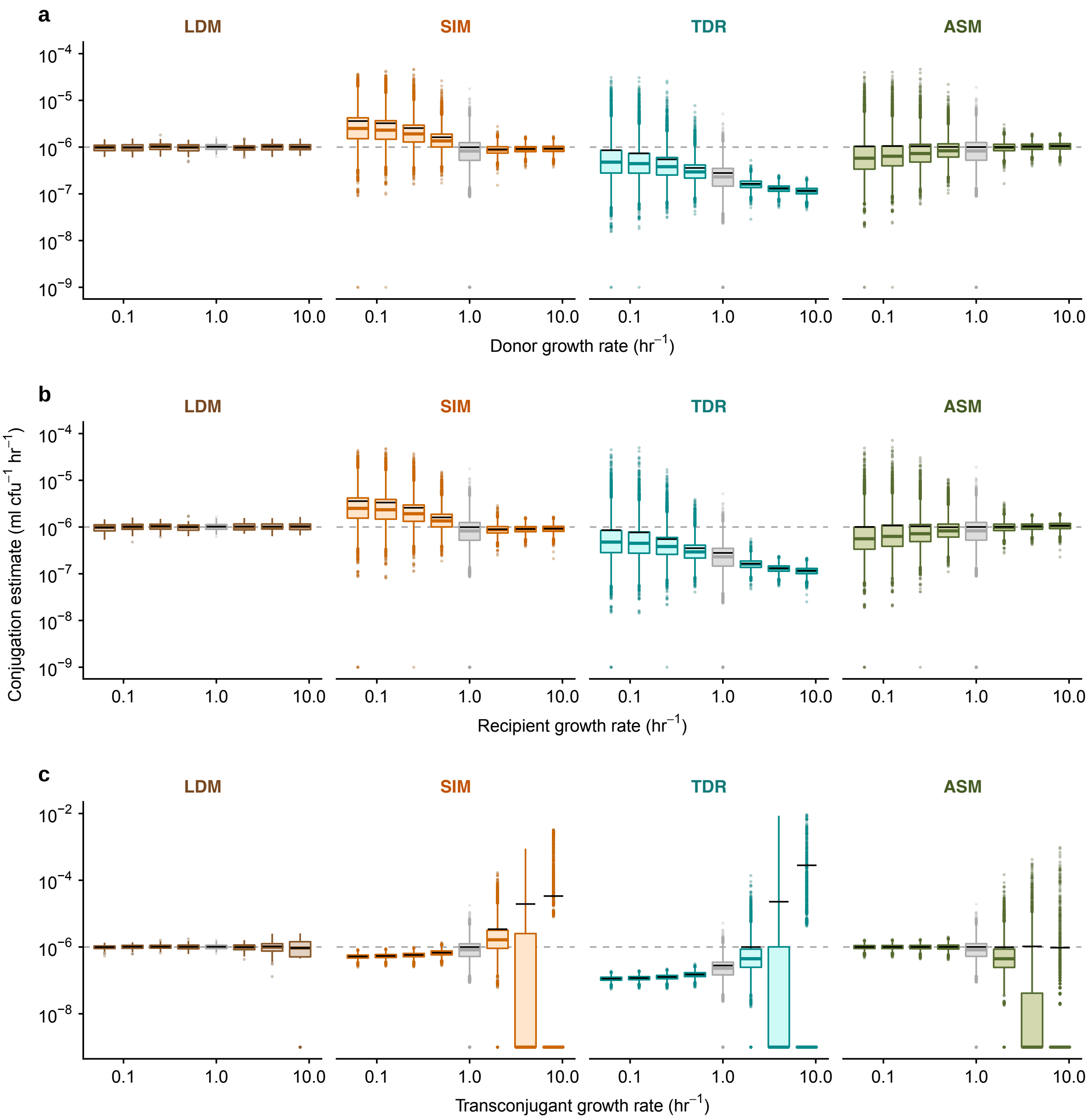


**SI Figure 2: The effect of population-specific heterogenous growth rate on estimating conjugation rate.** Boxplots are using the same graphical representation as SI Figure 1. (a, b, c) Unequal growth rates were explored over a range of growth rates for the donors, recipients, and transconjugants, respectively, namely $\psi_X \in \{0.0625, 0.125, 0.25, 0.5, 1, 2, 4, 8\}$. Zero estimates were set to $10^{-9}$ for plotting on a log axis.

*SI section 4c : The effect of unequal conjugation rates*

We expanded the analysis used in Figure 4b by calculating a conjugation rate estimate with two additional estimates, TDR and ASM (SI Figure 3). Like the conclusions drawn from Figure 4b with the effects of heterogenous conjugation rate, the LDM exhibited high accuracy and precision relative to other metrics. We note that we calculated the ASM metric in all scenarios and that in some cases the incubation time $\tilde{t}$ passed the critical time threshold ($t_{crit}$) where the ASM assumptions break down (see SI section 1a). The ASM estimate was included in all scenarios to be comprehensive and illustrate that implementing the assay after $t_{crit}$ can lead to bias. Given that the chosen incubation time $\tilde{t}$ to evaluate these simulations is early, it highlights that for a variety of parametric combinations proper implementation of the ASM metric is not possible.

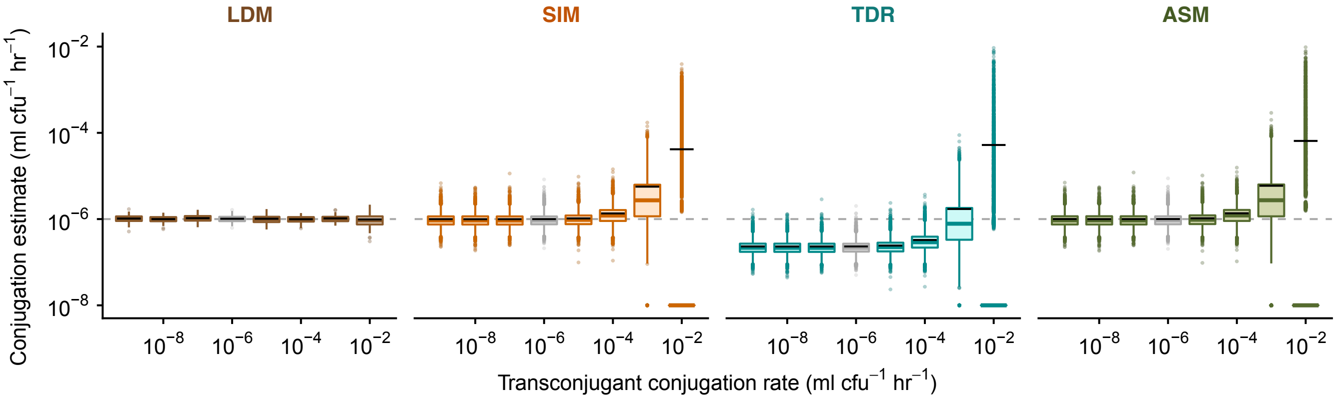


**SI Figure 3: The effect of heterogenous conjugation rate on estimating conjugation rate.** Boxplots are using the same graphical representation as SI Figure 1. Unequal conjugation rates were probed over a range of transconjugant conjugation rates, namely $\gamma_T \in \{10^{-9}, 10^{-8}, 10^{-7}, 10^{-6}, 10^{-5}, 10^{-4}, 10^{-3}, 10^{-2}\}$. Zero estimates were set to $10^{-8}$ for plotting on a log axis.

*SI section 4d : The effect of a non-zero plasmid loss rate*

We extended the base model (equations [1] - [3]) to include plasmid loss due to improper plasmid segregation. Thus, transconjugants are transformed into plasmid-free recipients due to improper segregation of the plasmid at rate $\tau_T$. The donors are transformed into plasmid-free cells due to improper segregation of the plasmid at rate $\tau_D$. Therefore, the extended model (equations [4.1] - [4.4]) tracks the change in density of a new population type, plasmid-free *former* donors ($F$). In total, the extended model describes the change in density of four populations ($D$, $R$, $T$, and $F$) due to various biological parameters: growth rates ($\psi_D$, $\psi_R$, $\psi_T$, and $\psi_F$), conjugation rates ($\gamma_{DR}$, $\gamma_{TR}$, $\gamma_{DF}$, and $\gamma_{TF}$), and plasmid loss rates ($\tau_D$ and $\tau_T$). Importantly, we note that all conjugation rates are dyad-specific (i.e., donor-recipient-specific); therefore, our simulation framework is built to allow all rates to be unique. Since the new population type are possible plasmid recipients, the subscript on the conjugation rate parameter now indicates the plasmid-bearing cell type and the plasmid-free cell type (e.g., $\gamma_{TF}$ indicates the conjugation rate between a transconjugant and a plasmid-free *former* donor).

$$\frac{dD}{dt} = \psi_D D + (\gamma_{DF} D + \gamma_{TF} T)F - \tau_D D, \qquad [4.1]$$

$$\frac{dR}{dt} = \psi_R R - (\gamma_{DR} D + \gamma_{TR} T)R + \tau_T T, \quad [4.2]$$

$$\frac{dT}{dt} = \psi_T T + (\gamma_{DR} D + \gamma_{TR} T)R - \tau_T T. \quad [4.3]$$

$$\frac{dF}{dt} = \psi_F F - (\gamma_{DF} D + \gamma_{TF} T)F + \tau_D D. \quad [4.4]$$

Plasmid loss due to improper segregation is a common occurrence in plasmid populations and violates a model assumption underlying all the conjugation rate estimates. We simulated a range of plasmid loss rates, ranging from low ($\tau_D$ = $\tau_T$ = $0.0001$) to high ($\tau_D$ = $\tau_T$ = $0.1$). The LDM had high accuracy and precision across all parameter settings (SI Figure 4). The effect of plasmid loss was undetectable even for an extremely high loss rate ($\tau_D$ = $\tau_T = 0.1$). Similarly, the effect of plasmid loss was undetectable on the other conjugation estimates compared to their performance with a zero loss rate. Thus, we find that all estimates appear robust with regards to an introduction of plasmid loss.

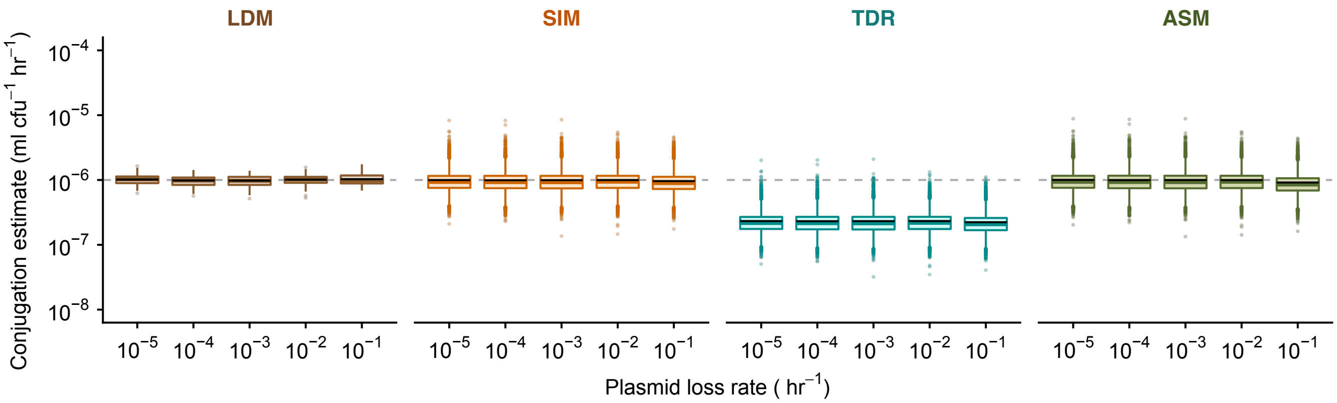


**SI Figure 4 : The effect of a non-zero plasmid loss rate on estimating conjugation rate.** Boxplots are using the same graphical representation as SI Figure 1. We explored improper plasmid segregation by considering a range of plasmid loss rates $\tau_D = \tau_T \in$ {0.00001, 0.0001, 0.001, 0.01, 0.1}.

*SI section 4e : The effect of incubation time using realistic parameter settings*

We expanded the analysis used in Figure 3 by calculating conjugation rate with two additional estimates, TDR and ASM. In addition, we explored the effects of incubation time in conjunction with other heterogenous parameter settings and a non-zero plasmid loss rate using realistic parameter settings. Given the computational expense of using realistic parameter values and higher initial densities, we explored five parameter combinations and the results are summarized in SI Figure 5. We set more reasonable initial densities of the donors and recipients ($D_0 = R_0 = 1 \times 10^5$) and a conjugation rate that is often reported in the literature ($\gamma_{DR} = \gamma_{DF} = \gamma_{TR} = \gamma_{TF} = 1 \times 10^{-14}$) unless otherwise indicated. The conjugation rate was estimated for each method at 30-minute intervals. For each time interval, we applied estimate-specific filters. For the LDM estimate, a 30-minute interval was shown if at least one parallel population had zero transconjugants. For the other estimates (SIM, TDR, and ASM), the 30-minute interval were shown if at least 90 percent of the simulated populations contained transconjugants at the incubation time. Even with only five parameter combinations, we note that the Gillespie algorithm is still computationally expensive when the densities get very large. Therefore, due to the longer incubation times needed for TDR, SIM, and ASM, only 100 populations of the 10,000 were simulated through the later time intervals; therefore, for each time point the box plots summarize 100 estimates for each method (TDR, SIM, ASM, and LDM).

The LDM estimate had high accuracy over all time points for all scenarios with precision increasing through time. The other estimates also become more precise over time. However, their greater precision over time was sometimes accompanied by decreased accuracy. We note these inaccuracies recaptured the qualitative patterns revealed in the parameter sweeps. Again, the LDM estimate performed as well or better than other estimates across incubation times.

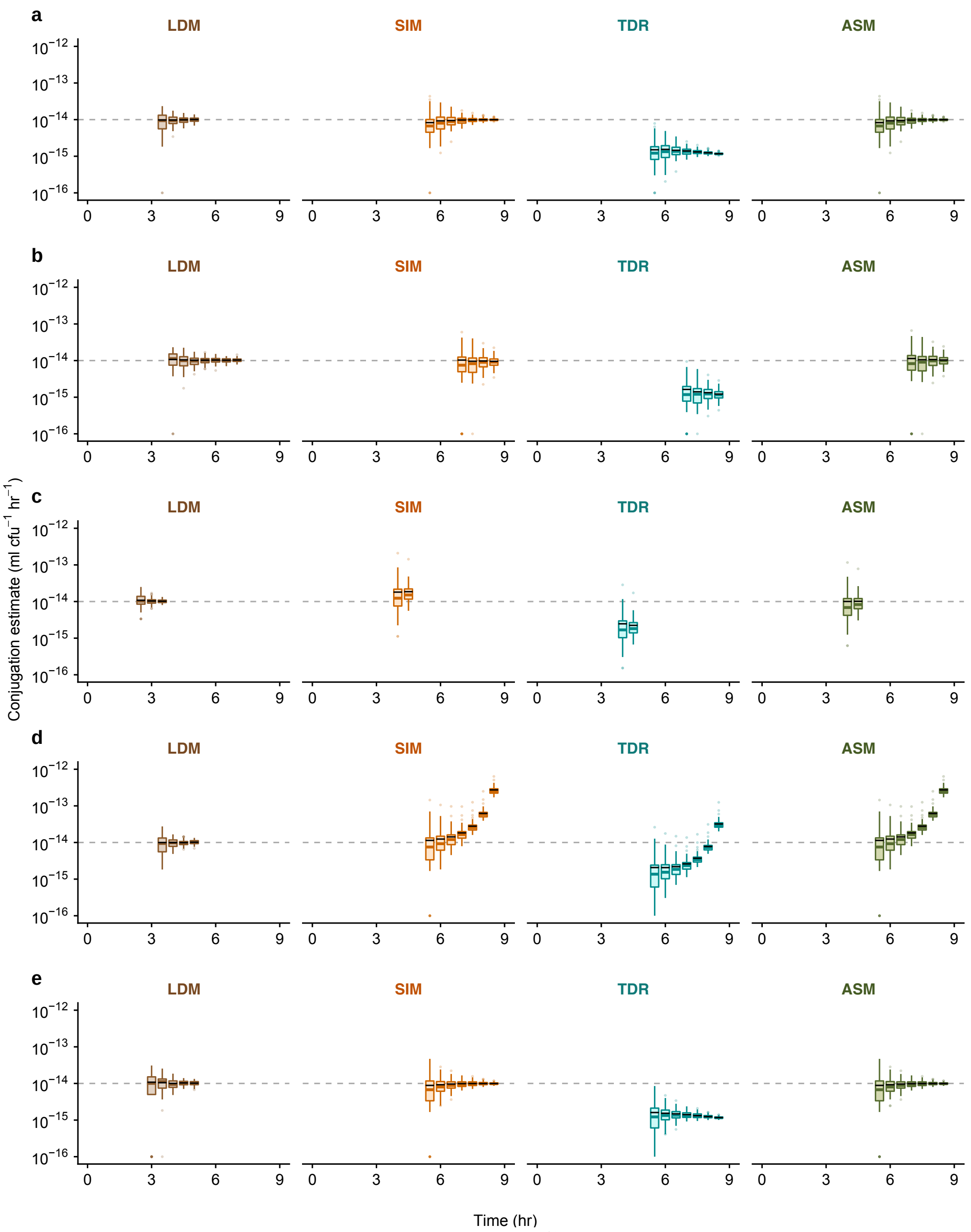


**SI Figure 5 : The effect of incubation time ($\tilde{t}$) on estimating conjugation rate.** The Gillespie algorithm was used to simulate population dynamics. Donor conjugation rate for each parameter combination was estimated at 30-minute intervals (summarized using boxplots with the same graphical convention as in Figure 3). The gray dashed line indicates the true value for the donor conjugation rate (here, $10^{-14}$). The baseline parameter values were $\psi_D = \psi_R = \psi_T = \psi_F = 1$ and $\gamma_{DR} = \gamma_{DF} = \gamma_{TR} = \gamma_{TF} = 1 \times 10^{-14}$. The dynamic variables were initialized with $D_0 = R_0 = 10^5$ and $T_0 = F_0 = 0$. The LDM, SIM, TDR, and ASM estimates are in separate plots with estimate specific colors (brown, orange, cyan, and green, respectively). (a) Baseline parameters were simulated as the non-heterogenous parameter comparison. (b) An unequal growth rate was simulated with $\psi_D = \psi_T = 0.5$. (c) An unequal growth rate was simulated with $\psi_R = \psi_T = 2$. (d) An unequal conjugation rate was simulated with $\gamma_{TR} = 10^{-8}$. (e) A non-zero plasmid loss rate was simulated with $\tau_D = \tau_T = 0.0001$.

*SI section 4f : Modified Levin et. al. model with Monod growth and conjugation*

Here we extend equations [4.1]-[4.4] to incorporate batch culture dynamics by tracking the change in resource concentration:

$$\frac{dD}{dt} = \psi_D(C)D + \gamma_{DF}(C)DF + \gamma_{TF}(C)TF - \tau_D(C)D, \quad [4.5]$$

$$\frac{dR}{dt} = \psi_R(C)R - \gamma_{DR}(C)DR - \gamma_{TR}(C)TR + \tau_T(C)T, \quad [4.6]$$

$$\frac{dT}{dt} = \psi_T(C)T + \gamma_{DR}(C)DR + \gamma_{TR}(C)TR - \tau_T(C)T, \quad [4.7]$$

$$\frac{dF}{dt} = \psi_F(C)F - \gamma_{DF}(C)DF - \gamma_{TF}(C)TF + \tau_D(C)D, \quad [4.8]$$

$$\frac{dC}{dt} = -(\psi_D(C)D + \psi_R(C)R + \psi_T(C)T + \psi_F(C)F)e. \quad [4.9]$$

where $e$ is the amount of resource required (µg) to produce a new cell. With the addition of a resource equation, there is an added assumption that growth, conjugation, and plasmid loss are Monod functions of resource concentration $C$:

$$\psi_X(C) = \psi_{X_{max}}\left(\frac{C}{Q+C}\right), \quad [4.10]$$

$$\gamma_{XY}(C) = \gamma_{XY_{max}}\left(\frac{C}{Q+C}\right), \quad [4.11]$$

$$\tau_X(C) = \tau_{X_{max}}\left(\frac{C}{Q+C}\right), \quad [4.12]$$

where $Q$ is the half saturation constant, and $\psi_{X_{max}}$, $\gamma_{XY_{max}}$, and $\tau_{X_{max}}$ are the maximum growth, conjugation, and plasmid loss rates for relevant cell types $X$ and $Y$, respectively. In other words, growth, conjugation, and plasmid loss decline and eventually turn off as resource concentration goes to zero.

*SI section 4g : Deterministic simulations with the Monod model using cross-species case study parameters*

We used equations [4.5]-[4.12] with the experimental parameters from the cross-species case study in Figure 6. Most of the parameters were from the average of six experiments ($D_0$ = 1.17 x $10^5$, $R_0$ = 3.33 x $10^4$, $\psi_D$ = 1.91, $\psi_R$ = 1.47, $\psi_T$ = 1.48, $\gamma_{DR}$ = 1.96 x $10^{-13}$, and $\gamma_{TR}$ = 1.96 x $10^{-7}$) with the remaining parameters informed by the 24 hour densities as to mimic the batch culture conditions of the experiment ($C_0$ = 4.41 x $10^9$, $Q$ = 1 x $10^7$, and $e$ = 1). We used the numerical solution to calculate the SIM estimate over time.

We compared the numerical solution to the actual experimental measurements from the cross-species experiments. The simulated density and conjugation estimate (SI Figure 6a solid lines) were similar to the average experimental densities and the experimental SIM estimate (SI Figure 6a circle data points). The LDM estimates for the cross-species conjugation rates ($\gamma_{DR}$ = 1.96 x $10^{-13}$) and the within-species ($\gamma_{TR}$ = 1.96 x $10^{-7}$) are sufficient to recapture a relatively inflated experimental SIM estimate. In contrast, homogenous conjugation rate with either the cross- or within-species conjugation rate does not closely align with the experimental data (SI Figure 6b and c, respectively).

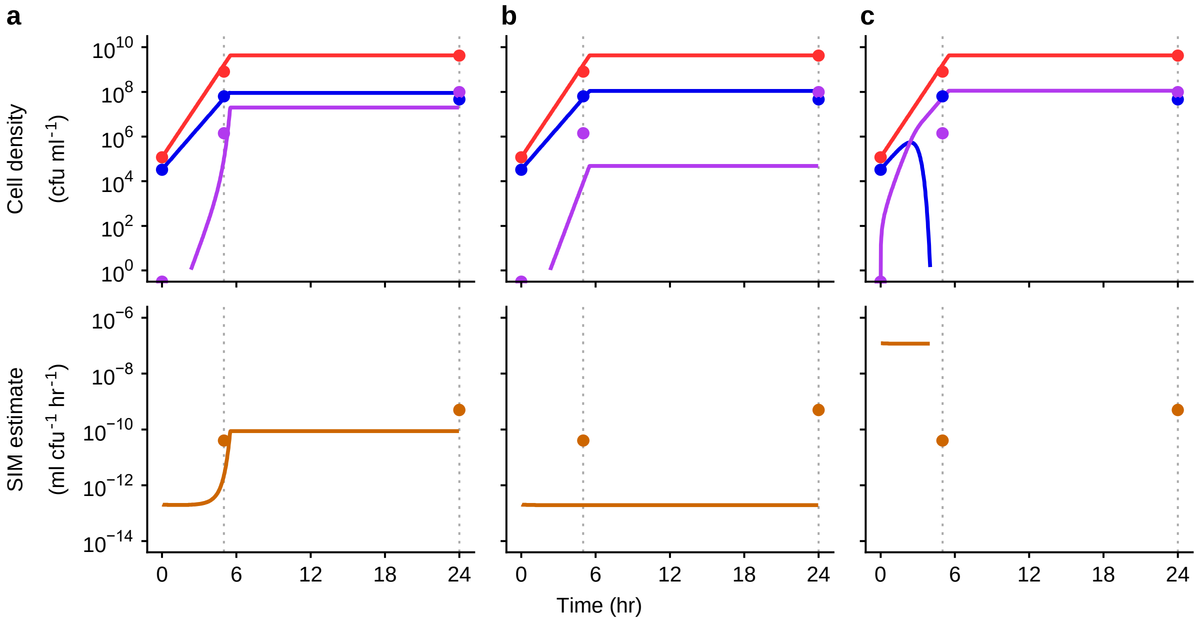


**SI Figure 6 : Numerical simulation of extended model with Monod functions using experimental parameters.** Deterministic numerical solutions of equations [4.5]-[4.12] showing donor, recipient, and transconjugant densities (red, blue, and purple solid trajectories, respectively) increasing over time using experimental parameter estimates ($D_0$ = 1.17 x $10^5$, $R_0$ = 3.33 x $10^4$, $\psi_D$ = 1.91, $\psi_R$ = 1.47, $\psi_T$ = 1.48, $\gamma_{DR}$ = 1.96 x $10^{-13}$, and $\gamma_{TR}$ = 1.96 x $10^{-7}$,) and batch culture parameters ($C_0$ = 4.41 x $10^9$, $Q$ = 1 x $10^7$, and $e$ = 1) unless otherwise indicated. The averaged experimental data is overlayed onto each part (circle data points) at both incubation times (grey dotted line at t = 5 and t = 24). (a) The numerical solution with the experimental parameter estimates were close to the experimental measurements. (b) A scenario with homogenous low conjugation rates ($\gamma_{DR}$ = $\gamma_{TR}$ = 1.96 x $10^{-13}$) deviates significantly from the experimental measurements. (c) A scenario with homogenous high conjugation rates ($\gamma_{DR}$ = $\gamma_{TR}$ = 1.96 x $10^{-7}$) deviates significantly from the experimental measurements.

*SI section 4h : Violation of the Levin et. al. model Monod equation assumptions*

We explored a model variation where the functional form of growth rate and conjugation rate are not proportional. While growth rates follow the Monod equation, conjugation rates are not dependent on resource and remain constant after resources are depleted. We found that this model results in a higher SIM estimate at 24 hours (SI Figure 7a) compared to the scenario where conjugation rate is proportional to growth rate (SI Figure 7b). It is worth noting that in this new model, the recipient pool is completely depleted which coincides with the SIM estimate hitting an asymptote. Note in SI Figure 7b, the SIM estimate also hits an asymptote; however, in this case it is due to the conjugation rates approaching zero as the resources are depleted.

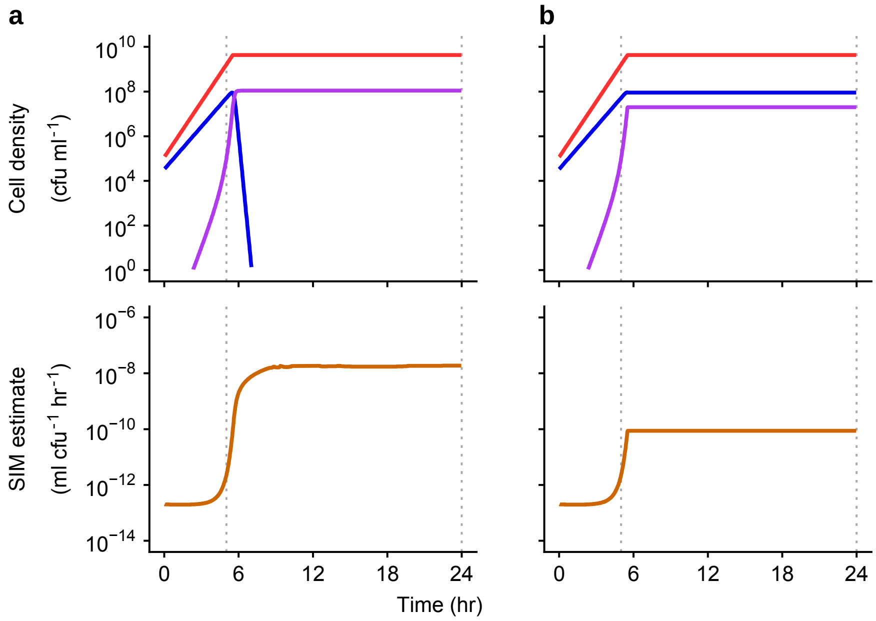


**SI Figure 7: Numerical simulation of modified model with a constant conjugation rate with Monod functions for growth.** The same equations and parameters from SI Figure 6 are used throughout unless otherwise indicated. (a) A model modification is made where conjugation rate is no longer proportional to growth rate. Specifically, conjugation rate is not resource dependent and is a constant. (b) The same panel in SI Figure 6a for comparison.

**SI section 5 : Experimental volume unit conversion using $f$**

In this section, we walk through the addition of $f$ to the LDM estimate. This is important to maintain the typical units $\mathrm{ml}/(\mathrm{h}\cdot\mathrm{cfu})$ used for reporting the conjugation rates. In the original differential equations [1]-[3], the units of the dynamic variables were $\mathrm{cfu}/\mathrm{ml}$. If we want to deal with numbers instead of density, the let us define a new volume unit termed the "$\mathrm{evu}$" standing for "experimental volume unit" where we will assume there are $f$ $\mathrm{evu}$'s per $\mathrm{ml}$. Focusing on the number of donors in the experiment (which we label $\breve{D}$), we have the following conversion:

$$\breve{D}\left(\frac{\mathrm{cfu}}{\mathrm{evu}}\right)=\frac{D\left(\frac{\mathrm{cfu}}{\mathrm{ml}}\right)}{f\,\frac{\mathrm{evu}}{\mathrm{ml}}},$$

Focusing on the numerical values (and ignoring the units for what follows), we have

$$\breve{D} = \frac{D}{f},$$

$$\breve{R} = \frac{R}{f},$$

$$\breve{T} = \frac{T}{f}.$$

In our original differential equations, let us multiply both sides of all the differential equations by $1/f$, yielding:

$$\frac{1}{f}\frac{dD}{dt} = \psi_D \frac{1}{f} D,$$

$$\frac{1}{f}\frac{dR}{dt} = \psi_R \frac{1}{f} R - \ (\gamma_D D + \gamma_T T)\frac{1}{f} R,$$

$$\frac{1}{f}\frac{dT}{dt} = \psi_T \frac{1}{f} T + \ \ (\gamma_D D + \gamma_T T)\frac{1}{f} R.$$

This can be reworked as

$$\frac{d\breve{D}}{dt} = \psi_D \breve{D},$$

$$\frac{d\breve{R}}{dt} = \psi_R \breve{R} - \ (\gamma_D D + \gamma_T T)\breve{R},$$

$$\frac{d\breve{T}}{dt} = \psi \breve{T} + (\gamma_D D + \gamma_T T)\breve{R}.$$

It follows that:

$$\frac{d\breve{D}}{dt} = \psi_D \breve{D},$$

$$\frac{d\breve{R}}{dt} = \psi_R \breve{R} - \ \left(f\gamma_D \breve{D} + f\gamma_T \breve{T}\right)\breve{R},$$

$$\frac{d\breve{T}}{dt} = \psi \breve{T} + \left(f\gamma_D \breve{D} + f\gamma_T \breve{T}\right)\breve{R}.$$

If we let

$$\breve{\gamma}_D = f\gamma_D,$$

and

$$\breve{\gamma}_T = f\gamma_T.$$

then the above system becomes

$$\frac{d\breve{D}}{dt} = \psi_D \breve{D},$$

$$\frac{d\breve{R}}{dt} = \psi_R \breve{R} - \ \left(\breve{\gamma}_D \breve{D} + \breve{\gamma}_T \breve{T}\right)\breve{R},$$

$$\frac{d\breve{T}}{dt} = \psi \breve{T} + \left(\breve{\gamma}_D \breve{D} + \breve{\gamma}_T \breve{T}\right)\breve{R}.$$

This set of equations tracks the number of cells (per $\mathrm{evu}$). Thus, if the above equations were used, then the derivations of the LDM estimate could flow exactly like we show in SI section 2. That is, the following will be correct:

$$\breve{\gamma}_D = -\ln p_0(\tilde{t})\left(\frac{\psi_D + \psi_R}{\breve{D}_0 \breve{R}_0\left(e^{(\psi_D+\psi_R)\tilde{t}} - 1\right)}\right)$$

Note, no correction is needed on $p_0(\tilde{t})$ as everything is in terms of numbers, which was how this quantity was derived. Because $\breve{D} = \frac{D}{f}$ and $\breve{R} = \frac{R}{f}$, we can rewrite the above as

$$\check{\gamma}_D = -\ln p_0(\tilde{t})\left(\frac{\psi_D+\psi_R}{\frac{D_0}{f}\frac{R_0}{f}\left(e^{(\psi_D+\psi_R)\tilde{t}}-1\right)}\right)$$

Or:

$$\frac{\check{\gamma}_D}{f} = f\left\{-\ln p_0(\tilde{t})\left(\frac{\psi_D+\psi_R}{D_0R_0\left(e^{(\psi_D+\psi_R)\tilde{t}}-1\right)}\right)\right\}$$

Because $\gamma_D = \frac{\check{\gamma}_D}{f}$, we have

$$\gamma_D = f\left\{-\ln p_0(\tilde{t})\left(\frac{\psi_D+\psi_R}{D_0R_0\left(e^{(\psi_D+\psi_R)\tilde{t}}-1\right)}\right)\right\}$$

Note that if our $\mathrm{evu}$ was 1 ml, then $f = 1$ and we could use our estimate exactly as written in equation [11]. Generally, we have to correct our original metric by multiplying by $f$.

### SI section 6 : Extended Experimental Methods and Results

#### *SI section 6a : Strains.*

*Escherichia coli* K-12 BW25113 from the Top Lab was used as the ancestor of the three *E. coli* strains in this study. To derive the first strain, *E. coli* BW25113 was grown overnight and plated onto LB agar supplemented with 100 µg ml$^{-1}$ streptomycin. A single streptomycin-resistant colony was selected and used to create an isogenic glycerol stock, *E. coli* K-12 BW25113 str$^R$, to be used as the plasmid-free *E. coli* recipient in this study (hereafter 'E(Ø)').

To derive the second strain, *E. coli* K-12 BW25113 was mixed with a host carrying the focal conjugative plasmid and incubated overnight in LB medium to facilitate plasmid transfer. The focal plasmid was the modified IncF conjugative plasmid F'42 (hereafter 'pF') in which a tetracycline resistance gene was inserted using lambda red recombination (8). The mixture was plated onto LB agar supplemented with 100 µg ml$^{-1}$ ampicillin (host selection) and 15 µg ml$^{-1}$ tetracycline (pF plasmid selection) to select for *E. coli* K-12 BW25113 host containing the pF plasmid. A single colony was selected and used to create an isogenic glycerol stock to be used as the plasmid-containing *E. coli* donor in this study (hereafter 'E(pF)').

To derive the third strain, E(pF) was mixed with E(Ø) and incubated overnight in growth medium to facilitate plasmid transfer. The mixture was plated onto LB agar supplemented with 100 µg ml$^{-1}$ streptomycin (host selection) and 15 µg ml$^{-1}$ tetracycline (plasmid selection) to select for *E. coli* K-12 BW25113 str$^R$ host containing the pF plasmid. A single colony was selected and used to create an isogenic glycerol stock to be used as a representative isogenic *E. coli* transconjugant in this study, hereafter '$E_T$(pF)' where the T subscript is added to distinguish this strain from the plasmid-bearing *E. coli* E(pF) strain, which is susceptible to streptomycin.

The *Klebsiella pneumoniae* strain Kp08 from Jordt *et. al.* (9) was used as the ancestor for the *K. pneumoniae* strain in this study. Kp08 was grown overnight and plated onto LB agar supplemented with 30 µg ml$^{-1}$ nalidixic acid. A single nalidixic-acid-resistant colony was selected and used to create an isogenic glycerol stock, *K. pneumoniae* Kp08 nal$^R$. Kp08 nal$^R$ was mixed with E(pF) and incubated overnight in growth medium to facilitate plasmid transfer. The mixture was plated onto LB agar supplemented with 30 µg ml$^{-1}$ nalidixic acid (host selection) and 15 µg ml$^{-1}$ tetracycline (plasmid selection) to select for *K.pneumoniae* Kp08 nal$^R$ host containing the pF plasmid. A single colony was selected and used to create an isogenic glycerol stock to be used as the plasmid-containing *K.*

*pneumoniae* donor in this study (hereafter 'K(pF)'). See SI Table 5 for a quick overview of the strains used in this study.

**SI Table 5: The strains used in this study.** Antibiotic abbreviations are as follows: tet = tetracycline, str = streptomycin, and nal = nalidixic acid, and the 'R' superscript indicates drug resistance in the strain.

| Strain | Host | Plasmid |
|---|---|---|
| E(pF) | *E. coli* K-12 BW25113 | F'42 $tet^R$ |
| K(pF) | *K. pneumoniae* Kp08 $nal^R$ | F'42 $tet^R$ |
| E(Ø) | *E. coli* K-12 BW25113 $str^R$ | None |
| $E_T$(pF) | *E. coli* K-12 BW25113 $str^R$ | F'42 $tet^R$ |

*SI section 6b : Growth rate assays.*

The strains (SI Table 5) were inoculated into LB medium from frozen glycerol stocks and grown overnight. The plasmid-containing cultures were supplemented with 15 µg $ml^{-1}$ tetracycline to select for maintenance of the plasmid. The saturated cultures were diluted 100-fold into LB medium to initiate a second 24 hours of growth (in order to acclimate the previously frozen strains to laboratory conditions). The acclimated cultures were then diluted 10,000-fold into LB growth medium and dispersed into 27 wells in a deep-well microtiter plate at a volume of 100 µl per well. Every hour, 30 µl was removed from three wells to determine cell density via selective plating (SI Figure 8a). The three replicate plates were averaged to estimate the cell density at each hour. The growth rates were calculated by taking the slope of each neighboring time point using equation [1.12] (SI Figure 8b). An incubation time was chosen for each strain to ensure bacterial cultures entered exponential growth before the start of the conjugation assay. An incubation time of 4 hours was used for both donors, E(pF) and K(pF), and the recipient, E(Ø).

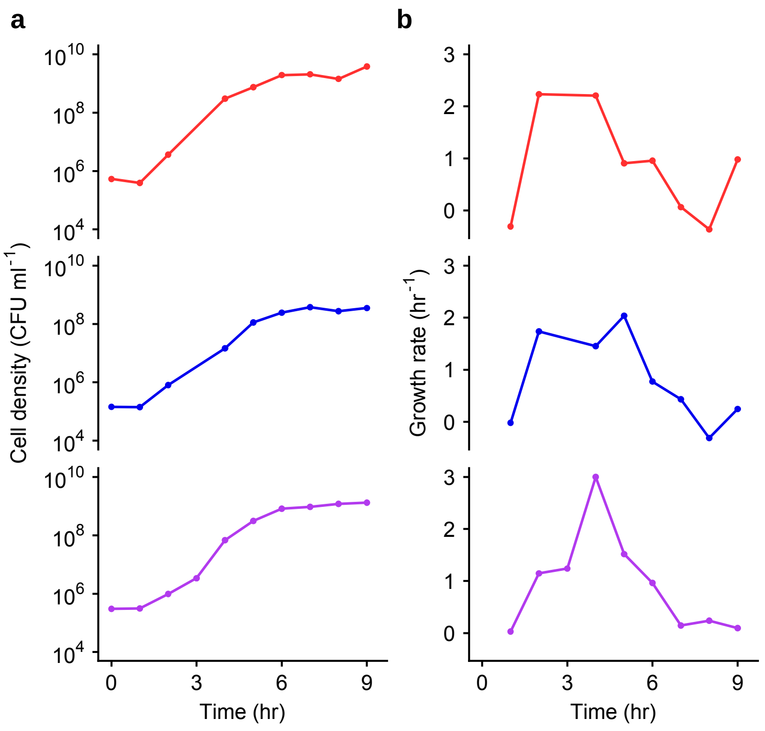


**SI Figure 8: The change in density and resulting growth rates of the relevant strains.** (a) Monocultures of K(pF), E(Ø), and E(pF) (red, blue, and, purple, respectively) were tracked over 9 hours of growth via plating. Bars indicate the standard error of the mean of three replicate cultures, but the standard error was so small in all cases that it is not visible in the plot. Note that at 3 hours a data point is missing for both K(pF) and E(Ø) due to plating error resulting in zero colonies and therefore no density estimate was available. (b) Using equation [1.12], the growth rates were calculated by taking the slope between each neighboring pair of time points (part a). Given the sampling frequency in the assay, the pair of time points were one hour apart except for the 4-hour data point where the 2-hour data point was used given the missing data. The growth rate estimate is plotted at the later time point used in the pair.

*SI section 6c : Minimum inhibitory concentration (MIC) assays.*

The strains (SI Table 5) were grown from glycerol stocks with two overnight incubations as previously described in SI section 6b. The acclimated cultures were diluted 100-fold into LB growth medium and dispersed into a column of wells in a deep-well plate at a volume of 500 µl per well. Then 500 µl of dual-antibiotic medium (streptomycin and tetracycline) was added to each well at increasing concentrations, forming a 2-fold gradient across the column. For each strain, this was repeated in three columns. After an overnight incubation, the well with the lowest concentration of the dual antibiotic medium across all replicates with no turbid growth was identified as the strain-specific MIC (SI Table 6). The concentration chosen for the transconjugant-selecting medium must be above the donor and recipient MIC, but below the transconjugant MIC. For this study, we proceeded with 7.5 µg ml$^{-1}$ tet + 25 µg ml$^{-1}$ str.

**SI Table 6: The dual-drug gradient MIC for the strains of interest.** The antibiotics used in the gradient were specific to the resistance profile of the transconjugant $E_T$(pF); streptomycin (str) and tetracycline (tet). The MIC data was used to identify the antibiotic concentration for the transconjugant-selecting medium used in both conjugation assays; cross- and within-species.

| Strain | Cell type | Str and tet gradient MIC |
|---|---|---|
| E(pF) | Donor | 1.88 μg ml$^{-1}$ tet + 6.25 μg ml$^{-1}$ str |
| K(pF) | Donor | 1.88 μg ml$^{-1}$ tet + 6.25 μg ml$^{-1}$ str |
| E(Ø) | Recipient | 3.75 μg ml$^{-1}$ tet + 12.5 μg ml$^{-1}$ str |
| $E_T$(pF) | Transconjugant | 15 μg ml$^{-1}$ tet + 50 μg ml$^{-1}$ str |

*SI section 6d : Extinction probability assays.*

A key component of the LDM conjugation protocol is differentiating parallel donor-recipient co-cultures that contain transconjugants from those that do not. This is done by adding transconjugant-selecting medium prepared at antibiotic concentrations below the MIC of the transconjugant and above the MIC of the donor and recipient. Given the low numbers of transconjugants in the co-cultures, the results from a recent study of Alexander and MacLean (10) have high relevance. First, the authors show that levels of antibiotic which select for the resistant cell type (i.e., below the MIC of the resistant strain) are sufficient to decrease the chance of outgrowth with very low cell numbers (e.g., a single cell). In the context of our current study, if the concentration of the transconjugant-selecting medium is too high then co-cultures that contain transconjugants could produce a non-turbid culture because the transconjugant cell(s) fail to establish a lineage. Therefore, to avoid spurious non-turbid wells in the LDM protocol, the probability that a transconjugant cell fails to establish (the transconjugant extinction probability) should ideally be 0 in the transconjugant-selecting medium. Second, the authors show that the presence of a sufficiently dense sensitive cell population in the environment can decrease the extinction probability of the resistant type. In the context of our current study, the presence of donors and recipients in the cultures may decrease the transconjugant extinction probability. Overall, a non-zero transconjugant extinction probability could lead to a biased estimate of the conjugation rate; therefore, it needs to be explicitly checked.

Inspired by the approach of Alexander and MacLean, we developed a similar approach to estimate the extinction probability of a transconjugant cell. First, we assume that a transconjugant cell has zero probability of extinction in antibiotic-free medium. While this assumption may be misplaced, it provides a starting point, and may itself be checked if there are reasons to doubt it holds. Second, we assume that a transconjugant cell has a specific probability of extinction in transconjugant-selecting medium with certain antibiotic concentrations given by the variable $x$, which is denoted $\pi_x$. Third, we assume that the lineage from every transconjugant cell in a population goes extinct independently. Consider a population of transconjugants distributed into many subpopulations containing transconjugant-selecting medium such that the average number of cells per subpopulations is initially $T$. Assuming an initial Poisson distribution, the fraction of subpopulations that leave zero transconjugant descendants, $P_x$ is:

$$P_x = \sum_{i=0}^{\infty} \frac{e^{-T}T^i}{i!}(\pi_x)^i = \frac{e^{-T}}{e^{-T\pi_x}} \sum_{i=0}^{\infty} \frac{e^{-T\pi_x}}{i!}(T\pi_x)^i = e^{-T(1-\pi_x)}.$$

By our assumption, when considering antibiotic-free medium, which we represent as $x = 0$, we have

$$P_0 = e^{-T(1-\pi_0)} = e^{-T}.$$

Thus, it is the case

$$T = -\ln P_0.$$

Given that

$$P_x = e^{\ln P_0 (1-\pi_x)},$$

we have a form to calculate the extinction probability in the transconjugant-selecting medium in the laboratory

$$\pi_x = 1 - \frac{\ln P_x}{\ln P_0}, \quad [6.1]$$

where $P_x$ is the fraction of non-turbid wells with transconjugant-selecting medium and $P_0$ is the fraction of non-turbid wells with antibiotic-free medium.

In the laboratory, we used the protocol implemented by Alexander and MacLean to estimate $\pi_x$ with a few adjustments. Briefly, the transconjugants were diluted (4 x $10^{-7}$) and 50 µl were dispersed into all wells in a deep-well microtiter plate. For the antibiotic-free condition, the wells were filled with LB medium to a final volume of 1 ml. For the transconjugant-selecting condition, the wells were filled with LB medium supplemented with transconjugant-selecting antibiotics (7.5 µg $ml^{-1}$ tet + 25 µg $ml^{-1}$ str, see SI section 6c for details) to a final volume of 1 ml. Both deep-well plates were incubated for 4 days. Using equation [6.1], we calculated a transconjugant extinction probability of 0.95 in the antibiotic concentration used for the transconjugant-selecting medium in this study.

Given that the extinction probability was non-negligible (i.e., $\pi_x > 0$), we ran a subsequent assay to estimate $\pi_x$ in the presence of sensitive cells (donors and recipients) at approximately the final densities that occur when the transconjugant-selecting medium is added for both mating assays (cross- and within-species, see SI Table 7) reported in this study. This provided a more accurate $\pi_x$ for correcting the LDM estimate (see SI section 7). In this experiment, the deep-well microtiter plates are prepared the same as above but supplemented with donor and recipient cells at the appropriate densities. As a result, we calculated mating-specific transconjugant extinction probabilities (SI Table 7). These mating-specific transconjugant extinction probabilities (given in SI Table 7) were used to correct the LDM estimate from each experimental replicate using equation [7.1].

Given the non-negligible extinction probability in the selective liquid medium, the extinction probability on the selective agar plates needed to be determined. We ran a subsequent assay to estimate $\pi_x$ for the donor-, recipient-, and transconjugant-selecting agar plates. Briefly, the monocultures of each strain were diluted ($10^{-5}$ and $10^{-6}$) and plated onto antibiotic-free plates and the appropriate selecting plates. We used a slightly altered form for calculating the agar extinction probability

$$\pi_x = 1 - \frac{C_x}{C_0}, \quad [6.2]$$

where $C_x$ is the number of colonies on the antibiotic-infused plate and $C_0$ is the number of colonies on the antibiotic-free plate for the same diluted culture. Using equation [6.2], we calculated each strain's extinction probability (see SI Table 8 for the antibiotic concentration used in the selective agar plates in this study). These strain-specific extinction probabilities were used to correct the density estimates from each experiment. We note that correcting the density estimates for the 24-hour data in 3 out of the 6 experiments resulted in negative estimates for the recipient density data. Specifically, the transconjugant density must be subtracted from the density calculated from the plate counts on agar supplemented with an antibiotic selecting for the recipient host, which also selects for transconjugants. Given the high transconjugant extinction probability on the transconjugant-selecting agar plates (see SI Table 8), the transconjugant density increases after the correction. Indeed, the transconjugant population becomes more common than the recipient population and the subtractive plating scheme becomes an unreliable approach to determine the recipient density (e.g., a "negative density" can result after correction). Since the exact recipient density cannot be determined due to its relative scarcity, if the recipient density went negative after subtraction, then the non-subtracted recipient density was used instead. An overestimate for recipient density leads to an underestimate for the SIM estimate at 24 hours; therefore, the differences between the cross-species LDM and SIM estimates shown in Figure 6 are conservative.

This section highlights the importance of non-zero extinction probabilities in selective conditions in the laboratory. Therefore, the extinction probabilities in selective liquid-medium and selective-agar plates need to be explicitly checked. If the extinction probabilities are indistinguishable from zero in each selective condition used, then the user can proceed, and no adjustments are necessary. However, a non-zero extinction probability is likely and can be a source of bias if not considered. We recommend two solutions. The first is to find a selection condition where the extinction probability is indistinguishable from zero. This option leans on the result from the Alexander and MacLean study which shows that the antibiotic concentration being sufficiently below the MIC of the focal strain can lower the extinction probability to a point that is indistinguishable from zero. We recognize that this solution may not be possible. For instance, the donor and recipient MIC for the transconjugant-selecting condition may be too close to the transconjugant MIC, such that there are no antibiotic concentrations that yield a zero transconjugant extinction probability and still counterselect donors and recipients. In this case, the user would proceed with the second solution where the extinction probabilities are used to compute a corrected estimate. This second solution was used in this study (see SI section 7).

**SI Table 7: Mating-specific transconjugant extinction probabilities with transconjugant-selecting liquid medium.** The donor and recipient densities were estimated using selective plating and were close to the final densities in the LDM conjugation protocol. Transconjugant-selective medium was prepared at the concentration used throughout the study (7.5 μg $ml^{-1}$ tet + 25 μg $ml^{-1}$ str).

| Mating | Donor density | Recipient density | $\boldsymbol{\pi_x}$ |
|---|---|---|---|
| within-species E(pF) to E(Ø) | $5 \times 10^4$ | $2 \times 10^4$ | 0.95 |
| cross-species K(pF) to E(Ø) | $1 \times 10^8$ | $7 \times 10^6$ | 0.93 |

**SI Table 8: Strain-specific extinction probabilities with selective-agar plates.** Donor-, recipient, and transconjugant-selective plates were prepared at concentrations that were used throughout the study (7.5 μg ml$^{-1}$ tet, 25 μg ml$^{-1}$ str, and 7.5 μg ml$^{-1}$ tet + 25 μg ml$^{-1}$ str, respectively).

| Strain | Selective-plate type | $\pi_x$ |
|---|---|---|
| E(pF) | Donor | 0.30 |
| K(pF) | Donor | 0.21 |
| E(Ø) | Recipient | 0.55 |
| $E_T$(pF) | Transconjugant | 0.99 |

*SI section 6e : Choosing an incubation time and initial density for executing the LDM conjugation assay.*

To find an incubation time and initial densities for executing the LDM protocol, all strains (SI Table 5) were prepared using the procedure in SI section 6b. We mixed exponentially growing donors and recipients in a large array of parallel co-cultures for a full factorial treatment of three initial densities and four incubation times (SI Figure 9a). We note that the resolution of initial densities and incubation times can be adjusted as needed. This is particularly useful if the conjugation rate is completely unknown. Alternatively, there could be good reasons for longer incubation times such as slow growth rates. For ease of explanation, we illustrate the protocol with a concrete example. Four columns were used for each initial density ($10^4$, $10^5$, and $10^6$-fold) where 2 rows were used for each incubation time (0, 1, 2, and 3 hours) resulting in 8 wells per density-time treatment. For each dilution, the exponentially growing donor and recipient cultures were diluted by the specific factor, mixed at equal volumes, and dispensed into the wells in the corresponding four columns at a volume of 100 μl per well (SI Figure 9a, black-bordered wells). At each incubation time, 900 μl of transconjugant-selecting medium (7.5 μg ml$^{-1}$ tetracycline and 25 μg ml$^{-1}$ streptomycin; see SI section 6c and SI section 6d) was added to each well in the corresponding two rows (SI Figure 9b, yellow-background). After the last time point ($t$ = 3 hours), the deep-well plate was incubated for 4 days. After the long incubation, we assessed the co-cultures within each time-density treatment for presence or absence of transconjugants by recording the turbidity (4 columns x 2 rows = 8 wells; SI Figure 9c). There were three outcomes possible for each time-density treatment: none of the co-cultures have transconjugants (gray-filled dot), all co-cultures have transconjugants (purple-filled dot), or there is both transconjugant-containing and transconjugant-free co-cultures (light-purple dot). The goal is to identify a density-time combination with the last outcome (i.e., both turbid and non-turbid co-cultures). These treatments meet the $\hat{p}_0(\tilde{t})$ condition (i.e., $0 < \hat{p}_0(\tilde{t}) < 1$). As a general expectation, a high donor conjugation rate ($\gamma_D$) will require shorter incubation times than a lower rate for a given initial density. For our matings (within- and cross-species), we found multiple density-time combinations that met the $\hat{p}_0(\tilde{t})$ condition. For the within-species mating assay, we chose a $10^3$-fold dilution and an incubation time of 1 hour and 15 minutes. For the cross-species mating assay, we chose the $10^3$-dilution and a 4-hour incubation time.

Even though multiple density-time treatments met the $\hat{p}_0(\tilde{t})$ condition, the final choice could not be made without the information from the controls. Thus, an additional deep-well plate was created (SI Figure 9d) to accompany the density-time plate containing the co-cultures (SI Figure 9a). This deep-well plate had the same factorial layout for densities (four columns) and incubation times (two rows) except the 8 wells within each treatment are not exclusively co-cultures. 3 of the wells contained monocultures of the three strains. Specifically, 100 µl of donor, recipient and transconjugant cultures were each placed in their own well (SI Figure 9d red-, blue- and purple-bordered wells). At a later point in the assay, these monocultures allowed us to determine that transconjugant-selecting medium prohibited growth of both donors and recipients, while permitting growth of transconjugants at each density-time treatment. An additional 2 wells contained monocultures of donors and recipients which are used to create a co-culture (in an empty well, dash-bordered well) during the assay itself at each incubation time (for each initial density). An additional 2 wells contained 100 µl of donor-recipient co-cultures which were used for selective plating to verify that the donors and recipients maintain a constant growth rate. At each incubation time, three events occurred in rapid succession. First, 30 µl was removed from each of the wells used to determine densities via selective plating. Second, donor and recipient monocultures were mixed at equal volumes into the empty well (SI Figure 9d, indicated by the gray arrows). Importantly, this well served as a control to verify that new transconjugant cells did not form via conjugation after transconjugant-selecting medium was added. Third, 900 µl of transconjugant-selecting medium was added to the first row of wells at the relevant time point (yellow background). The deep-well plate was incubated for 4 days. For the density-time combinations chosen, the control wells verified that the transconjugant-selecting medium operated as expected. In addition, the selective plating indicated the conditions under which the donors and recipients maintained constant growth.

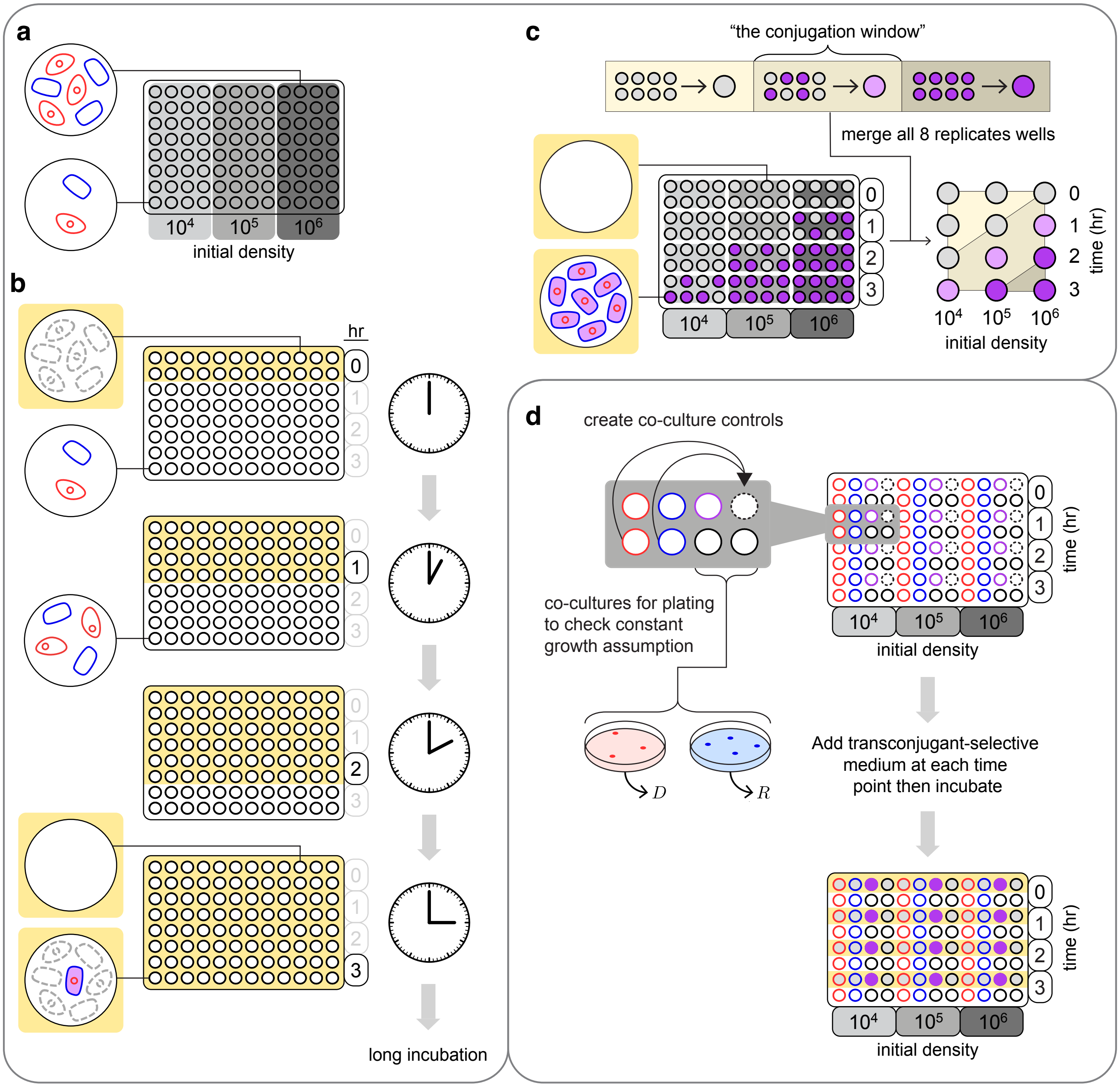


**SI Figure 9: Overview for finding an incubation time and initial densities for executing the LDM.** (a) the microtiter plate map designating the placement of the co-cultures over 10-fold increases in initial densities (different shades of gray). For simplicity, donors and recipients are at the same proportion in each co-culture. (b) Using the microtiter plate from part a, transconjugant-selecting medium (yellow-background) is added at each time designated by two rows in the microtiter plate. Two example wells from different density-time combinations are highlighted on the left. In the top example well, transconjugant-selecting medium is added immediately, inhibiting growth of donor and recipient cells (grey dashed cells), and resulting in a non-turbid well as no transconjugants formed. In the bottom example well, the donor and recipient population in the co-culture grow until transconjugant-selecting medium is added at 3-hours, inhibiting growth of donors and recipients, and permitting growth of the formed transconjugants. (c) After a lengthy incubation of the microtiter plate from part b, there are two well-types in the microtiter plate (bottom-left): transconjugant-containing (purple-filled) and transconjugant-free (gray-filled). For each density-time treatment, the 8 mating wells are considered as a group resulting in one of three outcomes (top): all transconjugant-free wells (gray dot), all transconjugant-containing wells (purple dot), a proportion of both well types (light-purple dot). Any treatment with a light-purple dot represents a viable combination of initial densities ($D_0'$ and $R_0'$) and incubation time ($\tilde{t}'$). (d) The microtiter plate with the control wells is set up with the same factorial layout used in part a except the 8 wells in each density-time treatment are not all co-cultures (black-bordered circles). Donor, recipient, and transconjugant monocultures serve as controls (red-, blue-, and purple-bordered wells, respectively). For the empty well (dash-bordered circles), donor and recipient monocultures are mixed into the empty well (indicated by grey arrows) to create a co-culture control at each time point to verify that diluting with transconjugant-selecting medium effectively prevents conjugation. In addition, the co-cultures are sampled at each time point to uncover densities and determine whether donors and recipients maintain constant growth. Subsequently, transconjugant-selecting medium is added to the microtiter plate at the same times as the microtiter plate in part a. The control wells inoculated with transconjugants should be turbid (purple-filled) while the monocultures with donors and recipients should be non-turbid. In addition, the co-cultures created at each time point for the different initial density treatments should be non-turbid.

### SI section 7 : Probability generating function, low-order moments, and failure to establish

Keller and Antal (11) studied a generalization of the process explored by Luria and Delbrück (12). To start, Keller and Antal consider a wildtype population expanding from a single cell as follows:

$$N_t = f(t) = e^{\delta t}.$$

Each wildtype cell generates a mutant cell at a rate $\nu'$, which grows as a stochastic birth process with rate $\alpha$ (Keller and Antal studied a supercritical birth-death process, but we will focus on the special case of a pure birth process). In this case, mutants form at a rate $\nu' f(t)$, such that the times of mutant arrival conform to a non-homogeneous Poisson process. We note that if we start with $N_0$ cells, then mutants form at a rate $N_0 \nu' f(t)$. Alternatively, we can set $\nu = N_0 \nu'$, such that mutants form at a rate $\nu f(t)$, which is the case explored by Keller and Antal.

Keller and Antal derive the probability generating function for the total number of mutants at an arbitrary time:

$$G(z,t) = \exp\left\{\frac{\nu}{\delta}\left(F\left(1,\kappa;1+\kappa;\frac{z}{z-1}e^{-\alpha t}\right) - e^{\delta t}F\left(1,\kappa;1+\kappa;\frac{z}{z-1}\right)\right)\right\},$$

where $F$ is the Gaussian hypergeometric function and $\kappa = \frac{\delta}{\alpha}$.

Our process of interest (the formation and growth of transconjugants) can be seen as an instance of their formulation by making the following substitutions:

$$\delta = \psi_D + \psi_R,$$
$$\nu = \gamma_D D_0 R_0,$$
$$\alpha = \psi_T.$$

With these substitutions, the generating function becomes:

$$G(z,t) = \exp\left\{\frac{\gamma_D D_0 R_0}{\psi_D + \psi_R}\left(F\left(1, \frac{\psi_D + \psi_R}{\psi_T}; 1 + \frac{\psi_D + \psi_R}{\psi_T}; \frac{z}{z-1}e^{-\psi_T t}\right) - e^{(\psi_D + \psi_R)t} F\left(1, \frac{\psi_D + \psi_R}{\psi_T}; 1 + \frac{\psi_D + \psi_R}{\psi_T}; \frac{z}{z-1}\right)\right)\right\}.$$

Because

$$G(z,t) = \sum_{n=0}^{\infty} p_n(t) z^n,$$

the probability of zero transconjugants now becomes straightforward (given $F\left(1, \frac{\psi_D+\psi_R}{\psi_T}; 1 + \frac{\psi_D+\psi_R}{\psi_T}; 0\right) = 1$):

$$p_0(t) = G(0,t) = \exp\left\{\frac{-\gamma_D D_0 R_0}{\psi_D + \psi_R}\left(e^{(\psi_D+\psi_R)t} - 1\right)\right\},$$

which agrees with the result from SI section 2.

Making the appropriate substitutions, we can also write the mean and variance (eqs. 8 and 9 from Keller and Antal) for the transconjugants:

$$E[T_t] = \begin{cases} \gamma_D D_0 R_0 e^{(\psi_D+\psi_R)t} t & \text{if } \psi_D + \psi_R = \psi_T \\ \dfrac{\gamma_D D_0 R_0\left(e^{(\psi_D+\psi_R)t} - e^{\psi_T t}\right)}{\psi_D + \psi_R - \psi_T} & \text{if } \psi_D + \psi_R \neq \psi_T \end{cases}$$

$$\mathrm{Var}[T_t] = \begin{cases} \dfrac{2\gamma_D D_0 R_0\left(e^{2(\psi_D+\psi_R)t} - e^{(\psi_D+\psi_R)t}\right)}{\psi_D + \psi_R} - \gamma_D D_0 R_0 e^{(\psi_D+\psi_R)t} t & \text{if } \psi_D + \psi_R = \psi_T \\ \dfrac{2\gamma_D D_0 R_0\left(e^{(\psi_D+\psi_R)t/2} - e^{(\psi_D+\psi_R)t}\right)}{\psi_D + \psi_R} + 2\gamma_D D_0 R_0 e^{(\psi_D+\psi_R)t} t & \text{if } \psi_D + \psi_R = 2\psi_T \\ \gamma_D D_0 R_0 \left\{\dfrac{2e^{2\psi_T t}(\psi_T - (\psi_D + \psi_R)) - e^{\psi_T t}(2\psi_T - (\psi_D + \psi_R)) + (\psi_D + \psi_R)e^{(\psi_D+\psi_R)t}}{(2\psi_T - (\psi_D + \psi_R))(\psi_T - (\psi_D + \psi_R))}\right\} & \text{otherwise} \end{cases}$$

We provide derivations for these expressions on the GitHub in Appendix VI. In all cases, the variance grows relative to the mean over time (see GitHub for Appendix VII for the derivations).

In our experiment, at time $\tilde{t}$, medium selecting for transconjugants is added to every mating culture. If every transconjugant always establishes a successful lineage, then every mating culture with one or more transconjugant cells at time $\tilde{t}$ will produce a turbid culture after a lengthy incubation. A more realistic scenario would be to assume that every transconjugant cell fails to establish a lineage with some probability, which we call $\pi$. If failure to establish occurs independently for each transconjugant, then the probability of a non-turbid culture after incubation ($P_{\mathrm{nt}}$) when selective medium was added at time $\tilde{t}$ is:

$$P_{\mathrm{nt}} = \sum_{n=0}^{\infty} p_n(\tilde{t}) \pi^n.$$

However, this is equivalent to an appropriate evaluation of the generating function:

$$P_{\mathrm{nt}} = G(\pi, \tilde{t}).$$

This can be rewritten as

$$P_{\mathrm{nt}} = \exp\left\{\frac{\gamma_D D_0 R_0}{\psi_D+\psi_R}\left(F\left(1,\frac{\psi_D+\psi_R}{\psi_T};1+\frac{\psi_D+\psi_R}{\psi_T};\frac{\pi}{\pi-1}e^{-\psi_T\tilde{t}}\right)\right.\right.$$
$$\left.\left.-\,e^{(\psi_D+\psi_R)\tilde{t}}F\left(1,\frac{\psi_D+\psi_R}{\psi_T};1+\frac{\psi_D+\psi_R}{\psi_T};\frac{\pi}{\pi-1}\right)\right)\right\}.$$

Solving for $\gamma_D$ yields

$$\gamma_D = \frac{-\ln\,(P_{\mathrm{nt}})(\psi_D+\psi_R)}{D_0R_0}\left(e^{(\psi_D+\psi_R)\tilde{t}}F\left(1,\frac{\psi_D+\psi_R}{\psi_T};1+\frac{\psi_D+\psi_R}{\psi_T};\frac{\pi}{\pi-1}\right)\right.$$
$$\left.-\,F\left(1,\frac{\psi_D+\psi_R}{\psi_T};1+\frac{\psi_D+\psi_R}{\psi_T};\frac{\pi}{\pi-1}e^{-\psi_T\tilde{t}}\right)\right)^{-1}$$

If the values of $D_0$ and $R_0$ are not the total initial numbers, but cell densities $(\mathrm{cfu/ml})$ in some volume for the mating culture (such that there are $f$ experimental volumes per ml) and we wish to measure conjugation rate in units $\mathrm{ml/(h\cdot cfu)}$, then must add a correction factor (see SI section 5), yielding

$$\gamma_D = f\,\frac{-\ln\,(P_{\mathrm{nt}})(\psi_D+\psi_R)}{D_0R_0}\left(e^{(\psi_D+\psi_R)\tilde{t}}F\left(1,\frac{\psi_D+\psi_R}{\psi_T};1+\frac{\psi_D+\psi_R}{\psi_T};\frac{\pi}{\pi-1}\right)\right.$$
$$\left.-\,F\left(1,\frac{\psi_D+\psi_R}{\psi_T};1+\frac{\psi_D+\psi_R}{\psi_T};\frac{\pi}{\pi-1}e^{-\psi_T\tilde{t}}\right)\right)^{-1} \quad [7.1]$$

First of all, we note that if every transconjugant establishes a lineage (i.e., $\pi=0$), then $P_{\mathrm{nt}} = p_0(\tilde{t})$ and equation [7.1] reduces to

$$\gamma_D = f\,\frac{-\ln\,(p_0(\tilde{t}))(\psi_D+\psi_R)}{D_0R_0\left(e^{(\psi_D+\psi_R)\tilde{t}}-1\right)},$$

which, using the maximum likelihood estimate for $p_0(\tilde{t})$, can be rewritten as

$$\gamma_D = f\left\{\frac{1}{\tilde{t}}[-\ln\hat{p}_0(\tilde{t})]\,\frac{\ln D_{\tilde{t}}R_{\tilde{t}}-\ln D_0R_0}{D_{\tilde{t}}R_{\tilde{t}}-D_0R_0}\right\},$$

and this is simply equation [13].

However, equation [7.1] is the more general expression. In SI section 6d, we discuss a method for estimating $\pi$. The maximum likelihood estimate for $P_{\mathrm{nt}}$ is the fraction of empty wells in the LDM protocol. Before, we called this $\hat{p}_0(\tilde{t})$, however, when there is positive probability that a transconjugant cell fails to establish (i.e., $\pi>0$), then generally $P_{\mathrm{nt}} > p_0(\tilde{t})$. Thus, we will denote the maximum likelihood estimate as $\hat{P}_{\mathrm{nt}}$ (the fraction of non-turbid wells).

If we let the density of transconjugants in a monoculture at times 0 and $\tilde{t}$ be $T_0^m$ and $T_{\tilde{t}}^m$, respectively (see SI section 6b) the following is the more general conjugation rate estimate (where all growth rates have been converted into estimated densities):

$$\gamma_D = f\,\frac{-\ln\,(\hat{P}_{\mathrm{nt}})\varsigma}{\tilde{t}}\left(D_{\tilde{t}}R_{\tilde{t}}F\left(1,\kappa;1+\kappa;\frac{\pi}{\pi-1}\right)\right.$$
$$\left.-\,D_0R_0F\left(1,\kappa;1+\kappa;\frac{\pi}{\pi-1}\frac{T_0^m}{T_{\tilde{t}}^m}\right)\right)^{-1} \quad [7.2]$$

with

$$\varsigma = \ln D_{\tilde{t}} R_{\tilde{t}} - \ln D_0 R_0,$$

and

$$\kappa = \frac{\varsigma}{\ln T_{\tilde{t}}^m - \ln T_0^m} = \frac{\ln D_{\tilde{t}} R_{\tilde{t}} - \ln D_0 R_0}{\ln T_{\tilde{t}}^m - \ln T_0^m}.$$

**SI section 8 : Variance in Estimates**

Here we will focus on two estimates, ASM and LDM, and ask about their variance (enabling us to compare precision). We will focus exclusively on the contributions to this variance coming from the stochasticity in the transconjugant numbers (i.e., ignoring contributions coming from assessment of initial and final donor and recipient populations). Details on some of the derivations in this section are given in Github Appendix VII.

We start with the ASM estimate (here we express the estimate in terms of growth rate parameters):

$$\gamma_D = \frac{\psi_D + \psi_R - \psi_T}{D_0 R_0 \left(e^{(\psi_D + \psi_R)\tilde{t}} - e^{\psi_T \tilde{t}}\right)} T_{\tilde{t}}.$$

Because we are only focusing on the contribution of the transconjugant variation, all parameters (initial densities and growth rates will be taken to be fixed). Thus, we can think about the ASM estimate as a random variable $\Gamma_{\mathrm{ASM}}$, where

$$\Gamma_{\mathrm{ASM}} = c_1 T_{\tilde{t}},$$

where the constant $c_1$ is

$$c_1 = \frac{\psi_D + \psi_R - \psi_T}{D_0 R_0 \left(e^{(\psi_D + \psi_R)\tilde{t}} - e^{\psi_T \tilde{t}}\right)}.$$

The variance of the ASM estimate is then

$$\mathrm{var}(\Gamma_{\mathrm{ASM}}) = {c_1}^2 \{\mathrm{var}(T_{\tilde{t}})\}.$$

But we have a closed form expression for $\mathrm{var}(T_{\tilde{t}})$. If $\psi_T \notin \{\psi_D + \psi_R, (\psi_D + \psi_R)/2\}$, we have

$$\mathrm{var}(\Gamma_{\mathrm{ASM}}) = \frac{\gamma_D(\psi_D + \psi_R - \psi_T)}{D_0 R_0} \left\{ \frac{(\psi_D + \psi_R) e^{(\psi_D + \psi_R)\tilde{t}} + (\psi_D + \psi_R - 2\psi_T) e^{\psi_T \tilde{t}} - (\psi_D + \psi_R - \psi_T) 2 e^{2\psi_T \tilde{t}}}{(\psi_D + \psi_R - 2\psi_T)\left(e^{(\psi_D + \psi_R)\tilde{t}} - e^{\psi_T \tilde{t}}\right)^2} \right\}.$$

The formulas for $\psi_T = \psi_D + \psi_R$ and $2\psi_T = \psi_D + \psi_R$ could also be derived via simple substitution (note, $\lim_{\psi_T \to \psi_D + \psi_R} c_1 = 1/(D_0 R_0 t e^{(\psi_D + \psi_R)\tilde{t}})$). These formulas allow us to project variance in the ASM estimate over time due to transconjugant variation if all parameters are known.

We now turn to the LDM estimate:

$$\gamma_D = -\ln \hat{p}_0(\tilde{t}) \left( \frac{\psi_D + \psi_R}{D_0 R_0 \left(e^{(\psi_D + \psi_R)\tilde{t}} - 1\right)} \right).$$

What we actually measure is the number of populations (or wells) that have no transconjugants (call this $w$) out of the total number of populations (or wells) tracked (call this W). As we show in Github Appendix IV, the maximum likelihood estimate of $p_0(\tilde{t})$ is

$$\hat{p}_0(\tilde{t}) = \frac{w}{\mathrm{W}}.$$

Of course, from experiment to experiment, there will be variance in the number of populations with no transconjugants. Let us consider a random variable $F$, which represents the fraction of total populations that have no transconjugants. The expectation of $F$ is (we drop the time argument for notational convenience):

$$\mathrm{E}[F] = p_0.$$

The second central moment of $F$ is

$$\mathrm{var}[F] = \frac{p_0(1-p_0)}{\mathrm{W}}.$$

Because we are only focusing on the contribution of the transconjugant variation, all parameters (initial densities and growth rates will be taken to be fixed). Thus, we can think about the LDM estimate as a random variable $\Gamma_{\mathrm{LDM}}$,

$$\Gamma_{\mathrm{LDM}} = c_2 \ln F,$$

where the constant $c_2$ is

$$c_2 = -\left(\frac{\psi_D + \psi_R}{D_0 R_0\left(e^{(\psi_D+\psi_R)\tilde{t}} - 1\right)}\right).$$

The variance of the LDM estimate is then

$$\mathrm{var}(\Gamma_{\mathrm{LDM}}) = {c_2}^2\{\mathrm{var}(\ln F)\}.$$

Here we use a first-order Taylor series approximation for $\ln F$ centered at $\mathrm{E}[F]$:

$$\ln F \approx \frac{F}{\mathrm{E}[F]} + \ln(\mathrm{E}[F]) - 1.$$

And we have

$$\mathrm{var}[\ln F] \approx \frac{1}{\mathrm{W}}\left(\frac{1}{p_0} - 1\right).$$

This approximation will be accurate when the deviation between $F$ and $\mathrm{E}[F]$ is very small (i.e., $\frac{|F-\mathrm{E}[F]|}{\mathrm{E}[F]} \ll 1$). As $\mathrm{W}$ (the number of replicate populations in the experiment) gets large, the distribution of $F$ will tighten around $\mathrm{E}[F]$, making the approximation more reasonable.

Now, we have the following expression for $p_0$ (reintroducing the time argument):

$$p_0(\tilde{t}) = \exp\left\{\frac{-\gamma_D D_0 R_0}{\psi_D + \psi_R}\left(e^{(\psi_D+\psi_R)\tilde{t}} - 1\right)\right\}.$$

Therefore,

$$\mathrm{var}[\ln F_{\tilde{t}}] \approx \frac{1}{\mathrm{W}}\left(\exp\left\{\frac{\gamma_D D_0 R_0}{\psi_D + \psi_R}\left(e^{(\psi_D+\psi_R)\tilde{t}} - 1\right)\right\} - 1\right),$$

where we make the time dependence of $F$ clear. Returning to the variance for the LDM estimate,

$$\mathrm{var}(\Gamma_{\mathrm{LDM}}) \approx \frac{1}{\mathrm{W}}\left(\frac{\psi_D + \psi_R}{D_0 R_0\left(e^{(\psi_D+\psi_R)\tilde{t}} - 1\right)}\right)^2\left(\exp\left\{\frac{\gamma_D D_0 R_0}{\psi_D + \psi_R}\left(e^{(\psi_D+\psi_R)\tilde{t}} - 1\right)\right\} - 1\right).$$

If we define

$$\xi_{\tilde{t}} = \frac{\psi_D + \psi_R}{D_0 R_0\left(e^{(\psi_D+\psi_R)\tilde{t}} - 1\right)},$$

then we have

$$\mathrm{var}(\Gamma_{\mathrm{LDM}}) \approx \frac{{\xi_{\tilde{t}}}^2}{\mathrm{W}}\left(e^{\left(\frac{\gamma_D}{\xi_{\tilde{t}}}\right)} - 1\right).$$

In SI Figure 10, we explore the variances (approximate in the case of LDM) as a function of time. The LDM estimates (for two different numbers of populations) are more precise (lower variance) for much of the time range. However, if the time gets too high ($\tilde{t} \approx 5$ for the parameter set shown in SI Figure 10), then the LDM variance blows up (while the ASM variance remains very low). In a case like this, the LDM is predicted to be more precise when the time of the assay is sufficiently low. In Appendix VII, we demonstrate this precision advantage for the LDM estimate mathematically.

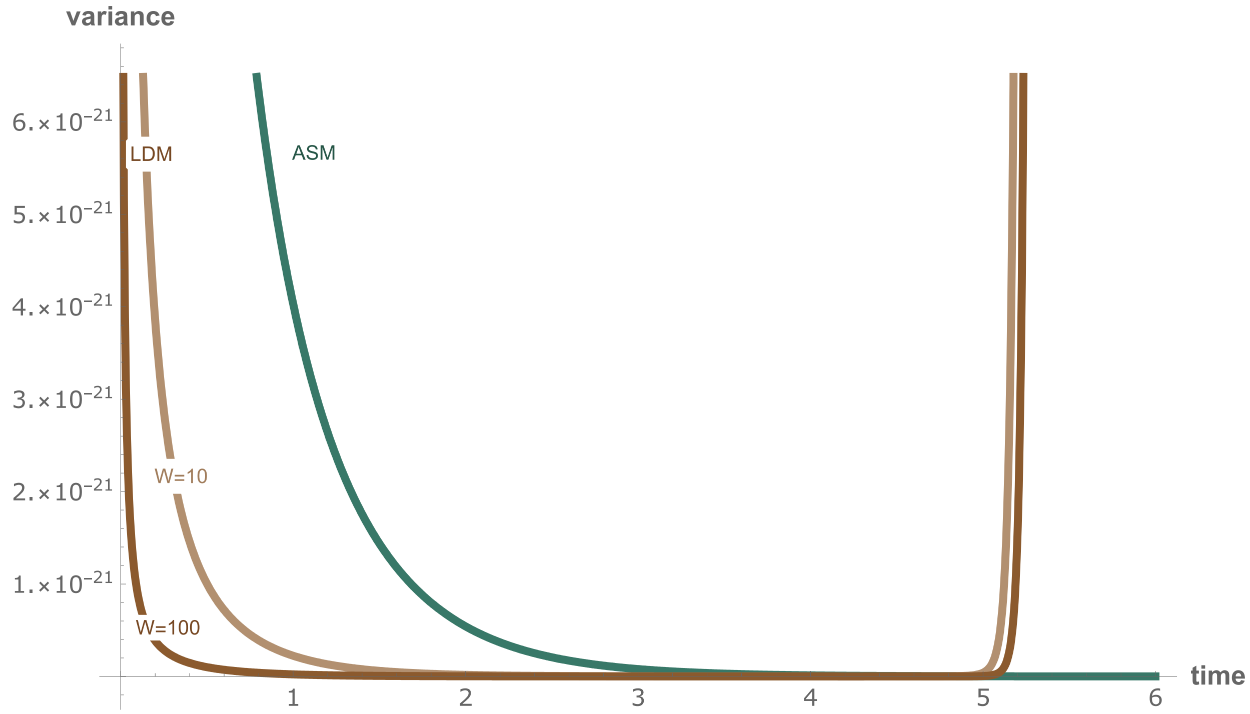


**SI Figure 10 : The variance of the ASM (green) and LDM (brown) estimates.** Different numbers of populations (W) are used for the LDM estimates, as indicated. The parameters used here are $\gamma_D = 10^{-12}$, $D_0 = R_0 = 10^4$, $\psi_D = 1$, and $\psi_R = \psi_T = 1.5$.

**References**


1. B. R. Levin, F. M. Stewart, V. A. Rice, The kinetics of conjugative plasmid transmission: fit of a simple mass action model. *Plasmid* **2**, 247–260 (1979).

2. X. Zhong, J. Droesch, R. Fox, E. M. Top, S. M. Krone, On the meaning and estimation of plasmid transfer rates for surface-associated and well-mixed bacterial populations. *J. Theor. Biol.* **294**, 144–152 (2012).

3. J. H. Bethke, *et al.*, Environmental and genetic determinants of plasmid mobility in pathogenic Escherichia coli. *Sci Adv* **6**, eaax3173 (2020).

4. A. J. Lopatkin, *et al.*, Antibiotics as a selective driver for conjugation dynamics. *Nat Microbiol* **1**, 16044 (2016).

5. L. Simonsen, D. M. Gordon, F. M. Stewart, B. R. Levin, Estimating the rate of plasmid transfer: an end-point method. *J. Gen. Microbiol.* **136**, 2319–2325 (1990).

6. T. A. Sysoeva, Y. Kim, J. Rodriguez, A. J. Lopatkin, Growth-stage-dependent regulation of conjugation. *AIChE* (2020).

7. J. S. Huisman, *et al.*, Estimating plasmid conjugation rates: a new computational tool and a critical comparison of methods. *bioRxiv*, 2020.03.09.980862 (2021).

8. H. L. Jordt, “Of E. coli and classrooms: Stories of persistence.” (2019).

9. H. Jordt, *et al.*, Coevolution of host-plasmid pairs facilitates the emergence of novel multidrug resistance. *Nat Ecol Evol* **4**, 863–869 (2020).

10. H. K. Alexander, R. C. MacLean, Stochastic bacterial population dynamics restrict the establishment of antibiotic resistance from single cells. *Proc. Natl. Acad. Sci. U. S. A.* **117**, 19455–19464 (2020).

11. P. Keller, T. Antal, Mutant number distribution in an exponentially growing population. *J. Stat. Mech.* **2015**, P01011 (2015).

12. S. E. Luria, M. Delbrück, Mutations of Bacteria from Virus Sensitivity to Virus Resistance. *Genetics* **28**, 491–511 (1943).
